# Operation of a TCA cycle subnetwork in the mammalian nucleus

**DOI:** 10.1101/2020.11.22.393413

**Authors:** Eleni Kafkia, Amparo Andres-Pons, Kerstin Ganter, Markus Seiler, Paula Jouhten, Filipa Pereira, Judith B Zaugg, Christophe Lancrin, Martin Beck, Kiran Raosaheb Patil

## Abstract

Nucleic acid and histone modifications critically depend on central metabolism for substrates and co-factors. Although a few enzymes related to the formation of these required metabolites have been reported in the nucleus, the corresponding metabolic pathways are considered to function elsewhere in the cell. Here we show that a substantial part of the mitochondrial tricarboxylic acid (TCA) cycle, the biosynthetic hub of epigenetic modification factors, is operational also in the nucleus. Using ^13^C-tracer analysis, we identified activity of glutamine-to-fumarate, citrate-to-succinate, and glutamine-to-aspartate routes in the nuclei of *HeLa* cells. Proximity labeling mass-spectrometry revealed a spatial vicinity of the involved enzymes with core nuclear proteins, supporting their nuclear location. We further show nuclear localization of aconitase 2 and 2-oxoglutarate dehydrogenase in mouse embryonic stem cells. Together, our results demonstrate operation of an extended metabolic pathway in the nucleus warranting a revision of the canonical view on metabolic compartmentalization and gene expression regulation.

---

Several of the chemical moieties and co-factors required for the covalent modifications of chromatin and RNA - acetyl-CoA, α-ketoglutarate, succinyl-CoA, 2-hydroxyglutarate, succinate and fumarate - originate in the tricarboxylic acid (TCA) cycle (*1–9*). Thus, beyond its canonical biosynthetic and bioenergetic role in cell function, the TCA cycle also plays a fundamental role in the spatiotemporal regulation of gene expression as well as in genome repair (*5, 6, 10–17*). The metabolites required for these modifications, or their direct precursors, are generally assumed to diffuse from mitochondria to the nuclear sites of need. This diffusion-centric scenario, however, contrasts with the emerging understanding of intracellular molecular crowding and phase separation (*18*), and disregards the reaction-diffusion case wherein the metabolite of interest can be *en route* sequestered by other enzymes. These considerations raise the possibility that some of the metabolites are produced inside the nucleus to ensure a timely supply to the corresponding nuclear processes. In support of this hypothesis, three individual TCA cycle enzymes (pyruvate dehydrogenase complex, PDC; α-ketoglutarate dehydrogenase complex, OGDC; and fumarate hydratase, FH) have been reported to be present in the nucleus (*14, 19–23*). Yet, these observations only partially alleviate the concerns against the diffusion-centric model. In particular, whether the precursors of these reactions, some of which are not highly abundant or stable in the cell, are diffusing into the nucleus or are in turn produced by other reactions occurring in the nucleus is unclear. Indeed, a nuclear pyruvate-to-α-ketoglutarate route was recently described, albeit during a specific stage of mouse embryonic development (*24*). Considering the problems with the diffusion-centric model, and the critical importance of multiple TCA cycle intermediates in chromatin and RNA modifications, we hypothesized that a large part of this metabolic network is operational in the mammalian nucleus.

To investigate the metabolic pathways operating in the nucleus, we incubated isolated nuclei from *HeLa* cells with uniformly ^13^C-labeled substrates representing key entry points or constituents of the TCA cycle (Fig. 1A, Methods). The nuclei were then washed to remove any extra-nuclear material to capture only the nucleoplasmic metabolite pools. In parallel, lysed cells were used as a control reflecting the enzymatic activities at the whole-cell level. The metabolic extracts were analyzed with gas chromatography - mass spectrometry (GC-MS) to quantify the relative abundance of the mass isotopomers of the TCA cycle intermediates.

**Fig. 1.**
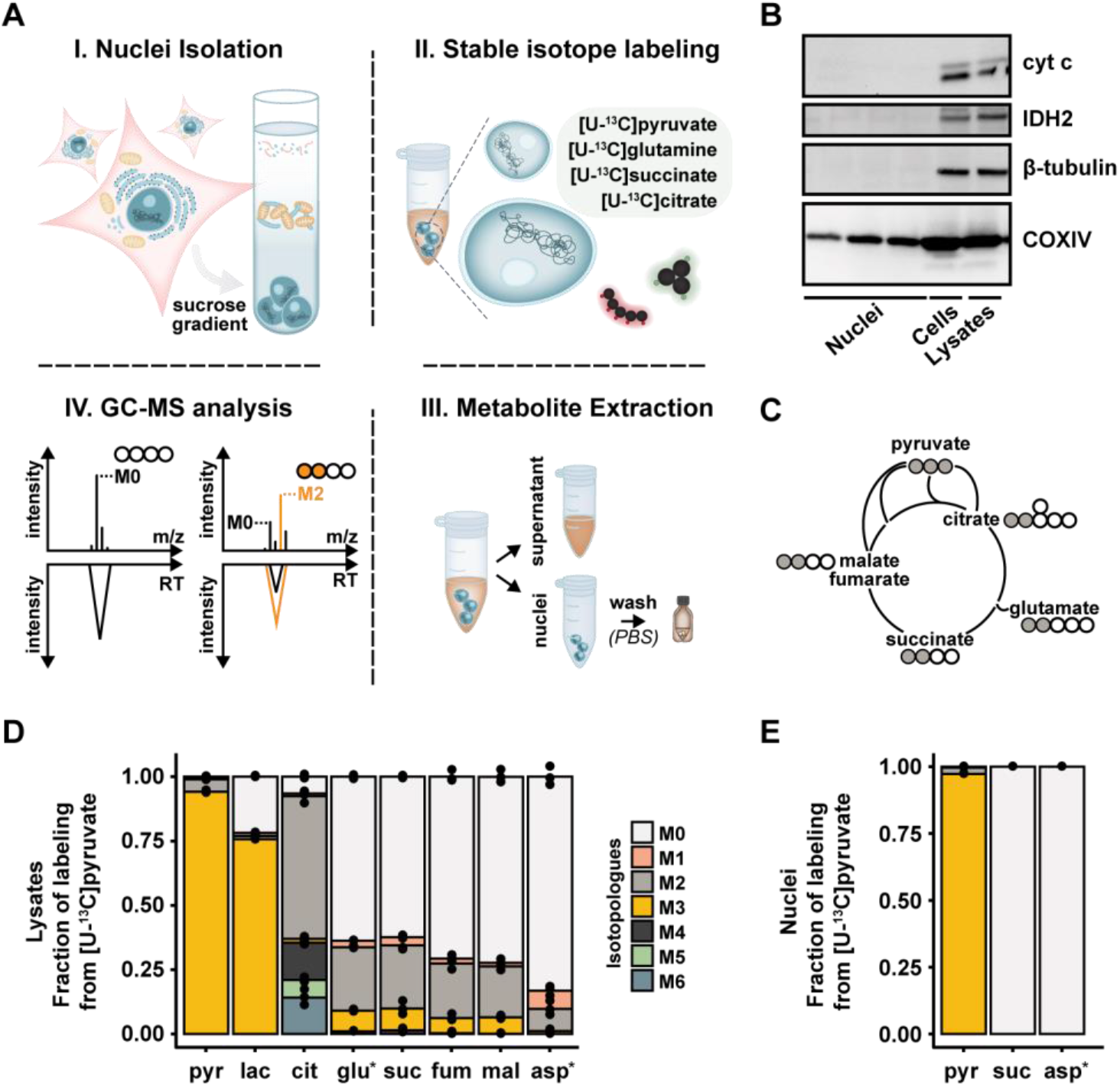
Probing the functional presence of multistep metabolic pathways in the nucleus. (**A**) Schematic overview of the ^13^C-labeling strategy in isolated nuclei. (**B**) Immunoblot analysis of isolated nuclei, cells and whole cell lysates for the mitochondrial shuttling protein cytochrome c (cyt c), the mitochondrial matrix protein isocitrate dehydrogenase 2 (IDH2), the cytoplasmic protein β-tubulin and the inner mitochondrial membrane protein COXIV. For the isolated nuclei, the data shown are from three independent nuclear isolations. For uncut membranes, see fig. S8 to S10. (**C**) Diagram showing the expected main mass isotopologue transitions of the TCA cycle intermediates in whole cell lysates incubated with [U-^13^C]pyruvate. Grey and empty circles represent ^13^C and ^12^C carbons, respectively. (**D** and **E**) Fraction of labeling of the different mass isotopologues (Mn; n, number of ^13^C-carbons) for each metabolite in whole cell lysates (**D**) and nuclei (**E**) incubated with [U-^13^C]pyruvate. Note that, for glutamate (glu*) and aspartate (asp*) the quantified ions correspond to a four and a three carbon fragment, respectively. For cell lysates, *n* = 3 biological replicates. For nuclei, *n* = 3 biological replicates. For all metabolomics data, see Tables S1 and S2. For panels D and E, data are presented as mean of the indicated biological replicates with individual data points shown. Abbreviations: pyr, pyruvate; lac, lactate; cit, citrate; glu, glutamate; suc, succinate; fum, fumarate; mal, malate; asp, aspartate.

To assess the purity of the nuclei isolations, we tested for the presence of the cytoplasmic protein β-tubulin, and two mitochondria-specific proteins, isocitrate dehydrogenase 2 and cytochrome c, representing mitochondrial matrix and trans-membrane shuttling activity, respectively. All three were not detected in the nuclear preparations (Fig. 1B). While the isolated nuclei appeared to carry fragments of mitochondrial membranes (COXIV, Fig. 1B), we verified by electron microscopy that the preparations were free of whole mitochondria (fig. S1). To quantitatively assay whether the mitochondrial membrane fragments interfered with the metabolite labelling, we contrasted the metabolic activities of the nuclei isolations and lysed cells following incubation with [U-^13^C]pyruvate. As expected, lysed cells were labeled in all measured TCA cycle metabolites (Fig. 1, C and D). Conversely, in isolated nuclei, the two detected TCA cycle intermediates, succinate and aspartate, did not incorporate any ^13^C in their carbon backbones (Fig. 1E). This result was consistent with previous studies in isolated nuclei from mammalian cells where pyruvate carbon entry in TCA metabolites other than acetyl-CoA was not observed (*19, 25*). Collectively, these control experiments mark the purity of the nuclei isolations regarding their enzymatic content.

In mitochondria, glutamine (/glutamate) is one of the main anaplerotic entry points feeding the TCA cycle, predominantly through α-ketoglutarate (*26, 27*). Considering that glutamine is among the most abundant amino acids in several tissues, in blood, and in common culture media (*28*), it is unlikely to be limited due to diffusion constraints. We therefore investigated whether it could supply the nucleus with the downstream metabolic intermediates of the TCA cycle. Tracing [U-^13^C]glutamine in isolated *HeLa* nuclei, we observed ^13^C enrichment in glutamate, succinate, fumarate and aspartate (Fig. 2A). We did not detect other TCA cycle intermediates in the nucleus, neither in labelled nor in unlabeled form. While the fractional labeling of glutamate and aspartate reached 90% already after 1h of incubation (fig. S2A), succinate and fumarate ^13^C fractions reached circa 45% after 5h (Fig. 2A and fig. S2A). The ^13^C fraction of succinate at 5h was smaller to that at 1h, consistent with its conversion to fumarate, the labeling of which showed a proportionate increase from 1h to 5h (fig. S2A). The control experiment with the lysed cells revealed additional metabolites being ^13^C-labeled, including malate and citrate, illustrating the activity of the whole TCA cycle as expected (Fig. 2B). To rule out the possibility that the absence of ^13^C-enriched intermediates downstream of fumarate in the nucleus was due to substrate concentration limitations, we incubated the nuclei (and lysed cells as a control) with [U-^13^C]succinate. While the lysed cells again showed the incorporation of ^13^C in fumarate, malate and citrate, in the nuclei only ^13^C-fumarate was detected (Fig. 2C) in agreement with the [U-^13^C]glutamine tracer experiment.

**Fig. 2.**
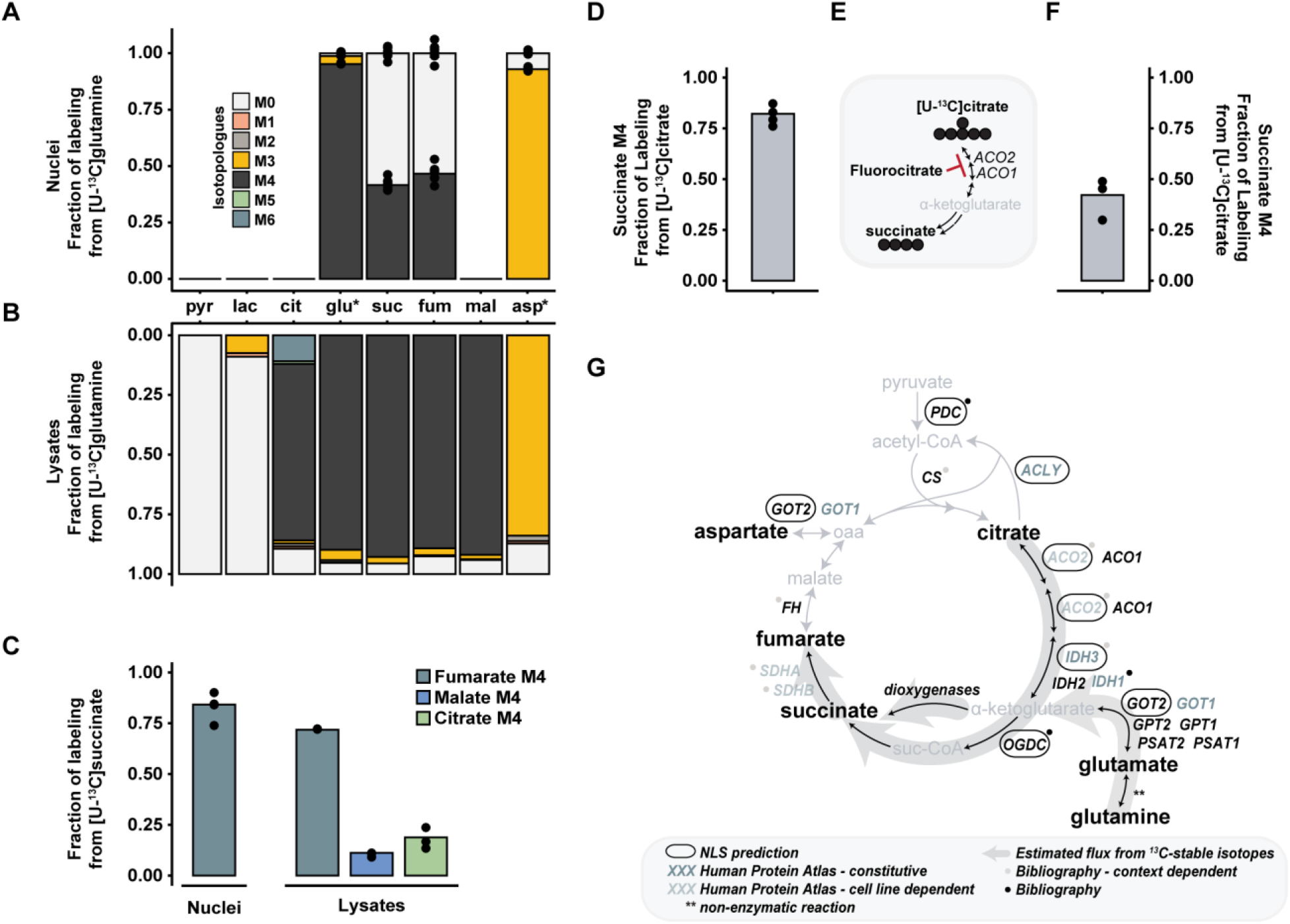
Glutamine and citrate reconstruct a subnetwork made of TCA cycle enzymes operating in the mammalian nucleus. (**A** and **B**) Fraction of labeling for the different mass isotopologues (Mn; n, number of ^13^C-carbons) for each metabolite in nuclei (**A**) and in whole cell lysates (**B**) incubated with [U-^13^C]glutamine. Note that, for glutamate (glu*) and aspartate (asp*) the quantified ions correspond to a four and a three carbon fragment, respectively. For nuclei, *n* = 6 biological replicates. For the whole cell lysates, one representative experiment is depicted. Abbreviations: pyr, pyruvate; lac, lactate; cit, citrate; glu, glutamate; suc, succinate; fum, fumarate; mal, malate; asp, aspartate. (**C**) Fraction of labeling for the M4 isotopologue of fumarate, malate and citrate in nuclei and whole cell lysates following incubation with [U-^13^C]succinate. For whole cell lysates, *n* = 3 biological replicates. For nuclei, *n* = 5 biological replicates. (**D**) Fraction of labeling for the M4 isotopologue of succinate in nuclei following incubation with [U-^13^C]citrate. *n* = 4 biological replicates. (**E**) Diagram of the potential initial enzymatic steps for the citrate-to-succinate pathway in the nucleus. (**F**) Effect of inhibition of aconitase activity by fluorocitrate on the fractional labeling of M4 isotopologue of succinate in isolated nuclei. *n* = 3 biological replicates. (**G**) Schematic of the traditional TCA cycle in mitochondria overlaid with a summary of the ^13^C-labeling flux estimations in isolated nuclei (light gray background arrows), the nuclear localization signal (NLS) predictions (black box around the enzymes), the nuclear immunofluorescence evidence from the Human Protein Atlas (HPA) (green and light green letters denote a constitutive or cell-line dependent nuclear presence, respectively) and bibliography (black and light gray dots denote a constitutive or context-dependent nuclear presence, respectively). For panels A to F, data are presented as mean of the indicated biological replicates with individual data points shown. For all metabolomics data, see also Tables S1 and S2.

To assess the utilization of carbon sources other than glutamine in the nucleus, we next focused on citrate, a key metabolic intermediate known to locally provide acetyl-CoA for chromatin modifications (*15, 29*). Incubating *HeLa* nuclei with [U-^13^C]citrate resulted in the detection and ^13^C labeling of succinate (Fig. 2D). The absence of labelling in fumarate, in nuclei as well as in the lysed cells (fig. S2B), was likely due to the known properties of citrate as metal chelator and allosteric inhibitor of several metabolic enzymes (*30*). To ascertain the enzymatic nature of the citrate to succinate conversion, we used fluorocitrate, an inhibitor of aconitase that catalyzes the first reaction in this route (Fig. 2E). The presence of fluorocitrate led to a two-fold reduction in the ^13^C fractional labeling of succinate (Fig. 2F), corroborating that aconitase is active in the *HeLa* nucleus.

The results from the glutamine and citrate labelling experiments support these as sources of TCA cycle intermediates in the *HeLa* nucleus. To more broadly examine the nuclear presence of the corresponding enzymes, we next explored the Human Protein Atlas images (*31*). We observed that aconitase 2 (ACO2) and isocitrate dehydrogenase 3 (IDH3) that catalyze the citrate-to-α-ketoglutarate reactions, exhibit nuclear localization that is either cell-line dependent (ACO2) or ubiquitous (IDH3G) across all examined cells (Fig. 2G). Additional TCA cycle enzymes, as well as enzymes that catalyze equivalent reactions elsewhere in the cell, likewise were found to exhibit signal for nuclear localization. These included isocitrate dehydrogenase 1 (IDH1), succinate dehydrogenase subunits A (SDHA) and B (SDHB), and various aminotransferases (e.g. GOT1) responsible for glutamate carbon entry to TCA cycle through α-ketoglutarate formation (Fig. 2G). Mutant IDH1 and the heterodimer SDHA-SDHB have been previously reported in the nucleus, albeit with no functional relevance (*32, 33*). Further, published proteomics data from isolated nuclei included several TCA cycle components (*34*). We therefore searched the sequences of TCA cycle enzyme for nuclear localization signals (NLS). Interestingly, we found putative canonical nuclear localization signals (NLS) for every enzymatic step, save citrate synthase (CS), from pyruvate up to the generation of succinyl-CoA, (Fig. 2G). Collectively, these independent and orthogonal evidences - Human Protein Atlas, proteomics, and NLS - provide a localization support to every enzymatic step implicated by our functional labeling results.

To further attest the nuclear localization of the implicated metabolic route, we next aimed to identify interacting and proximal proteins for a subset of TCA cycle enzymes using *in vivo* proximity-dependent biotinylation (BioID) coupled to mass spectrometry (*35*). The selected bait enzymes covered part of the citrate to succinate axis: ACO2, IDH3G subunit from IDH3 complex, IDH1, and OGDH subunit from OGDC complex. As a negative control, we used IDH2, a mitochondrial enzyme with no evidence for nuclear localization. We also included, despite having no evidence of downstream TCA cycle products in our labelling assays, pyruvate dehydrogenase B (PDHB) as a potential nuclear localized enzyme based on previous reports (*19, 21*).

Gene ontology (GO) enrichment analysis of the results from the BioID assay for these five enzymes affirmed that the vast majority of the biotinylated proteins (hereafter termed putative interactors) resided in the primary location of the corresponding bait enzyme; the mitochondria for all, except IDH1 which mainly localizes to cytoplasm (Fig. 3A, clusters 2 and 3, Tables S3 and S4). Yet, we identified a group of putative interactors, shared by OGDH and IDH3G, featuring the nucleolus as the top enriched subcellular compartment (Fig. 3A, cluster 1, Tables S3 and S4). This cluster did not show any interactions with PDHB and IDH2, consistent with the sole mitochondrial localization of the latter (Tables S3 and S4). GO analysis also highlighted activities closely linked with nucleolus, notably RNA binding and metabolic processing of rRNAs (Fig. 3A and Table S4). This is in agreement with recent studies reporting TCA cycle enzymes as RNA binding proteins (*36, 37*). Taken together, these data, while attesting the localization of the bait enzymes to their primary compartment, also support their additional nuclear presence, particularly discernible for OGDH and IDH3G.

**Fig. 3.**
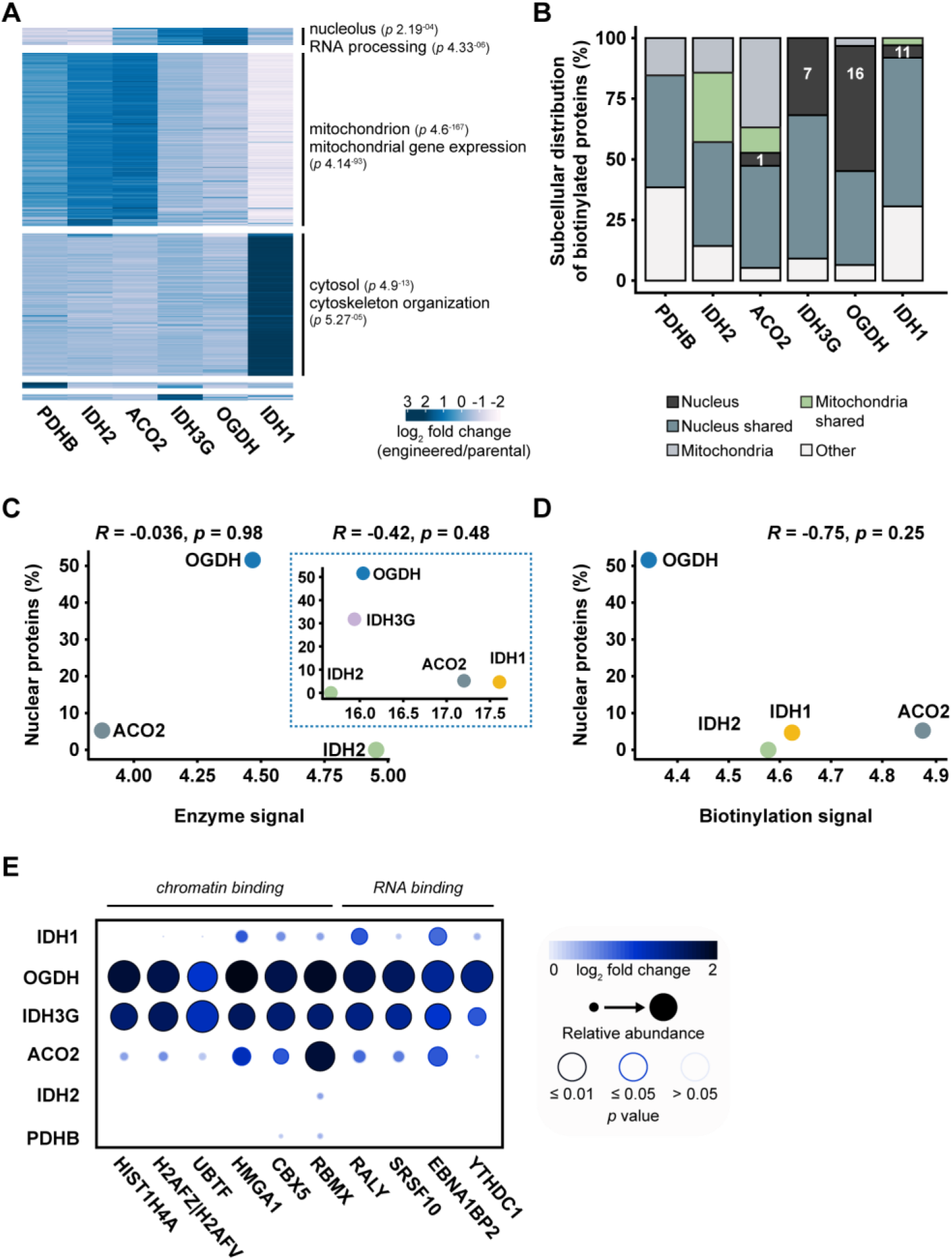
Proximity biotinylation mass spectrometry reveals core nuclear proteins as putative interacting partners for IDH1, IDH3G, OGDH and ACO2. (**A**) Heat map, clustering and gene ontology enrichment (GO) analysis for the significant putative interactors associated with each bait enzyme. The color gradient reflects the averaged log2 fold change of the abundance of each putative interacting protein in the cells expressing a bait enzyme relative to the parental *HeLa* cells. Hierarchical clustering was based on the “Euclidean” distance metric using the “ward.D2” linkage. Each cluster was analyzed for GO enrichment with g:Profiler; for the first three clusters, the most significant term for “Cellular Component” (upper) and “Biological Process” (lower) is depicted. See also Methods and Tables S3 and S4. (**B**) Subcellular distribution (percentage) of the putative interactors associated with each bait enzyme. See also Methods and Table S5. (**C**) Pearson correlation analysis of the percentage of nuclear proteins with the abundance of selected bait enzymes as defined by immunofluorescence (main panel) or by proteomics (insert panel). Immunofluorescence was performed using antibodies against the indicated endogenous enzymes. Proteomics data were obtained from (*38*). See also fig. S3A. (**D**) Pearson correlation analysis of the percentage of nuclear proteins with the biotinylation levels of selected cells expressing the bait enzymes. Biotinylation corresponds to the putative interacting partners and was defined by immunofluorescence using fluorescently-labeled streptavidin. See also fig. S3, B and C. (**E**) Dotplot of selected putative interacting proteins associated with different bait enzymes. Input data represent the averaged log2 fold change in the abundance and respective *p* values of each putative interactor in the cells expressing a bait enzyme relative to the parental *HeLa* cells. See also Methods, Tables S3 and S5.

To more stringently assess the localization of the interactors of the TCA cycle enzymes, we next identified the top putative interactors for each of the bait enzymes by contrasting against IDH2 hits (negative control, mitochondrial) (Methods). Inspection of the known localizations of the resulting putative interactors revealed a fraction characterized as exclusively nuclear for OGDH, IDH3G, IDH1 and ACO2 (Fig. 3B, Table S6). Importantly, no nuclei-only localized putative interactors were detected for the mitochondria-exclusive IDH2 (as compared against ACO2). None of the PDHB interactors also show exclusive nuclear localization, despite its reported nuclear presence, which can be attributed to the cell cycle phase-dependent localization to the nucleus (*19*). We also checked whether the capture of nuclear proteins could be attributed to high abundance of the bait proteins suggestive of noise or artefacts. No such bias was observed, regardless of whether the abundances were estimated by using immunofluorescence, mass-spectrometry (*38*) (Fig. 3C and fig. S3A), or the whole cell biotinylation levels (Fig. 3D and fig. S3B). Amongst the putative nuclear interaction partners, in addition to RNA binding proteins, we identified chromatin binding and remodeling factors (e.g. RBMX, HMGA1) and specific histones (e.g. HIST1H4A, H2AFZ) (Fig. 3E, Table S6). The latter is in accord with the recently reported interaction between OGDC and acetyltransferase 2A mediating the succinylation of histone lysine residues (*22*).

We next visualize the subcellular distribution of selected bait enzymes and their corresponding putative interactors, using immunofluorescence microscopy. The enzymes were detected in their expected primary compartment, while IDH3G was additionally present in the region of nucleolus (fig. S4). The staining pattern of the putative interactors (corresponding to the biotinylation signal as assessed with fluorescently-labeled streptavidin), followed the primary localization of the respective bait enzymes, while a discernible nuclear signal was detected only in the case of IDH1 (Fig. 4A and fig. S5). To minimize the more abundant mitochondrial staining that could obscure the nuclear signal in all other cases for which the nuclear localization was suggested by our labelling and BioID assays (i.e. OGDH, IDH3G and ACO2), we examined the staining for the putative interacting proteins (biotinylation) in isolated nuclei. For the case of IDH2, the nuclear biotinylation levels were similar to those of cells not expressing a bait enzyme (Fig. 4B and fig. S6, B and C), attesting the sole mitochondrial location. In comparison, in all other cases, there was a significantly higher degree of nuclear biotinylation (p value < 0.01) (Fig. 4B and fig. S6, B and C). Together with the BioID mass-spectrometry results, the microscopically visible putative interacting partners in the region of the nucleus for IDH1, ACO2, OGDH and IDH3G further support their nuclear presence.

**Fig. 4.**
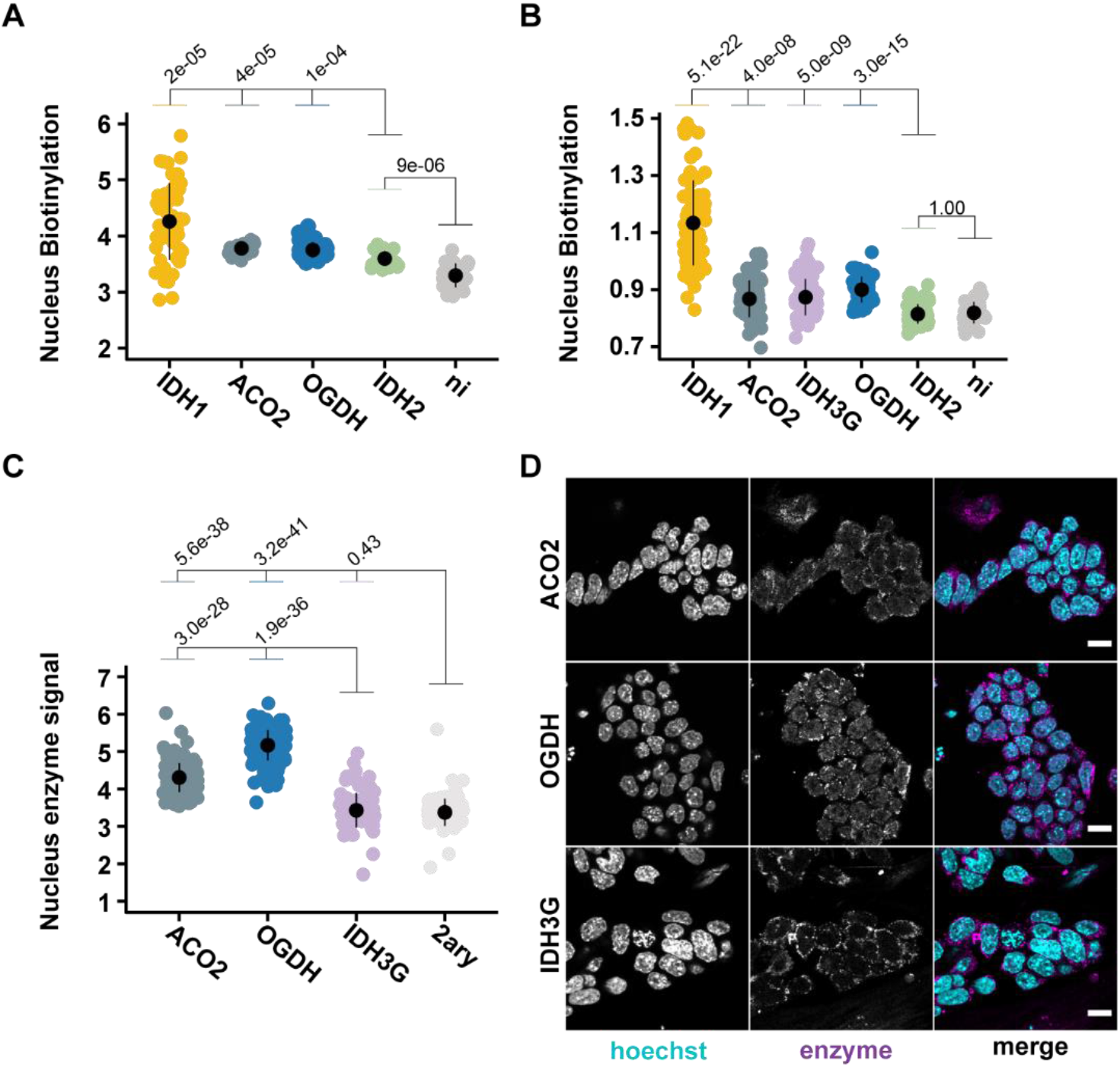
The subcellular distribution of the metabolic enzymes and the putative interacting partners reveal the existence of a nuclear neighborhood. (**A** and **B**) Assessment of the signal for the putative interactors in the nuclear region of whole cells expressing the bait enzymes (A) or in isolated nuclei from cell lines expressing the bait enzymes (B). The putative interactors correspond to the biotinylation signal as defined by immunofluorescence using fluorescently-labeled streptavidin. “ni” refers to cells where the expression of the bait enzyme was not induced with tetracycline; in the particular case we used the cell line of OGDH. For panel A, from left to right, *n* = 52, 14, 40, 28 and 34 quantified cells. For panel B, from left to right, *n* = 73, 75, 69, 55, 63 and 37 quantified nuclei. See also fig. S5 and S6. (**C**) Quantification of ACO2, OGDH and IDH3G in the nucleus of embryonic stem cells. The quantification was based on immunofluorescence using antibodies against the indicated endogenous enzymes. “2ary” refers to cells stained only with the secondary, fluorophore-conjugated, antibody. From left to right, *n* = 183, 166, 132 and 133 quantified cells. See also fig. S7. (**E**) Representative immunofluorescence images of embryonic stem cells stained for ACO2, OGDH and IDH3G (magenta). DNA was stained with Hoechst (cyan). Scale bars, 15µm. See also fig. S7. In (A) to (C), points represent quantification (log2 scale) of individual cells or nuclei, with the population mean and standard deviation calculated for each case. In (B), the biotinylation levels were normalized to Hoechst levels. In (A) to (C), significance was assessed for the indicated comparisons using the Wilcoxon rank sum test.

The visibility of the putative interactors in isolated nuclei but not in whole cells is in line with the expected low abundance of the enzymes in the nucleus relative to their primary location. We hypothesized that this situation would be different in developing cells wherein extensive chromatin remodeling depends on the availability of TCA cycle metabolites (*5, 6*). We therefore examined the presence of IDH3G, ACO2 and OGDH in mouse embryonic stem cells by immunofluorescence and detected a clear nuclear signal for the latter two enzymes (Fig. 4, C and D, and fig. S7). This provides a hitherto unrecognized spatial context for the link between metabolism and the epigenetic modulations that underline the differentiation and renewal of embryonic stem cells.

Taken together, our results show that not only specific enzymatic steps but more extended TCA cycle subnetworks operate in the mammalian nuclei. These nuclear pathways utilize citrate and glutamine/glutamate as source molecules and produce downstream metabolites critical for modifications on chromatin and RNA. Proximity of the involved enzymes with canonical nuclear proteins with roles in histones and nucleic acid modifications is in accord with the ‘on-site’ production of critical metabolites. This interpretation is further strengthened by the fact that cells under developmental context exhibited a more prominent nuclear presence of key enzymes than *HeLa* cells. Our data challenges the notion confining the TCA cycle activity to mitochondria, and brings forward the distribution of the metabolic activity between mitochondria and nucleus as a fundamental aspect of gene expression regulation.

## Supporting information

Tables_S1-S10_kafkia_et_al

## Acknowledgements

We acknowledge support from the following at the European Molecular Biology Laboratory (EMBL, Heidelberg, Germany): P. Ronchi from the Electron Microscopy Core Facility (EMCF), M. Rettel and F. Stein from the Proteomics Core Facility (PCF), S. Schnorrenberg, M. Lampe and S. Terjung from the Advanced Light Microscopy Facility (ALMF). We also thank M. T. Mackmull and M. Beck’s laboratory for help establishing the nuclear isolation and proximity biotinylation protocols, and providing the pcDNA5-pDEST-BioID-FLAG-N-term plasmid, the pcDNA5-pDEST-BioID-FLAG-C-term plasmid, the *HeLa* Kyoto cell line and the Flp-In T-REx *HeLa* cell line.

## Author contributions

E.K. and K.R.P. conceived the project, designed experiments, analyzed data and wrote the manuscript. E.K. performed most of the experiments and data analysis with assistance as follows: A.A.P. performed the immunoblotting analysis, K.G. performed the immunofluorescence staining and image acquisition of embryonic stem cells, M.S. performed the nuclear localization signal analysis, P.J. set up the natural abundance correction of the ^13^C-labeling results, and F.P. helped with the design of constructs for BioID assays. J.B.Z. helped in data interpretation. C.L. contributed to and oversaw the embryonic stem cells experiments. M.B. contributed to and oversaw the nuclei isolation and the proximity biotinylation experiments. K.R.P. oversaw the project.

## Data availability

The mass spectrometry proteomics data have been deposited to the ProteomeXchange Consortium (http://proteomecentral.proteomexchange.org) via the PRIDE (*60*) partner repository with the dataset identifier PXD022608. The metabolomics data have been deposited in MetaboLights (*61*) with the database identifier MTBLS2235. All original imaging data are available upon request.

## Materials and Methods

### Nuclei Isolation

*HeLa* Kyoto cells (RRID: CVCL_1922) were maintained under standard cultivation conditions (37°C, 5% CO2) in Dulbecco’s modified Eagle medium (DMEM, high glucose, GlutaMAX, Thermo Fisher Scientific, 61965026) supplemented with 10% heat-inactivated fetal bovine serum (FBS, Thermo Fisher Scientific, 26140079). For nuclei experiments, the cells were seeded in 245×245 mm plates (2.0 × 10^6^ cells/plate) and cultivated for 3 days. The nuclei isolation method was adapted from Ori and colleagues (*39*). The below described quantities refer to the collection of one 245 mm plate. Briefly, cells were washed three times with 10 mL PBS followed by addition of Trypsin-EDTA (Thermo Fisher Scientific, 25300096). Once cells were detached from the plate, 20 mL of cultivation medium was added and the cells were collected. From here onwards, all steps were performed in ice-cold conditions. Following centrifugation at 500g for 5 minutes, the cell pellet was washed with 10 mL PBS and centrifuged under the same conditions. Subsequently, the cell pellet was resuspended in 7.5 mL hypotonic buffer A (50 mM Tris-HCl pH 7.5, 1 µg/ mL aprotinin (Carl-Roth, A162.1) and 0.5 µg/ mL leupeptin (Carl-Roth, CN33.3)) and incubated on ice for 30 minutes. Once the cells were swollen, rapture was achieved utilizing a Dounce homogenizer with pestle B. Cell lysis efficiency was monitored through light microscopy. Once cells were sufficiently lysed, the hypotonic buffer A was adjusted to buffer B (0.25 M sucrose, 50 mM Tris-HCl pH 7.5, 25 mM KCl, 5 mM MgCl2, 2 mM DTT, 1 µg/mL aprotinin and 0.5 µg/mL leupeptin). A fraction of whole cell lysates was immediately used for the 13C-labeling experiments. The rest of the lysates were centrifuged at 1000g for 8 minutes and the pelleted nuclei were resuspended in 10 mL buffer B. Following centrifugation, the nuclei pellet was resuspended in 3 mL buffer consisted of one part buffer B and two parts buffer C (2.3M sucrose, 50 mM Tris-HCl pH7.5, 25 mM KCl, 5 mM MgCl2, 2 mM DTT, 1 µg/mL aprotinin, 0.5 µg/mL leupeptin). The resuspended nuclei were transferred to ultracentrifuge tubes (Beckman, #3440057) and 1.5 mL of buffer C was slowly placed at the bottom of the tubes with a needle of at least 18G. The nuclei were centrifuged at 21000 g for 30 minutes in a Beckman ultracentrifuge equipped with a SW55Ti rotor. Subsequently, the interphase and the supernatant were carefully aspirated via vacuum, and the nuclei pellet was resuspended in 1 mL buffer B and centrifuged at 1000g for 8 minutes. This step was repeated two more times. The nuclei were immediately utilized for the 13C-labeling experiments. Alternatively, samples were saved at −80°C for immunoblot analysis.

### 13C-labeling experiments and metabolite extraction

Freshly isolated nuclei were resuspended in 500 μL incubation buffer containing 0.25 M sucrose, 50 mM Tris-HCl pH 7.5, 25 mM KCl, 5 mM MgCl2, 2 mM DTT, 1 µg/mL aprotinin, 0.5 µg/mL leupeptin, 1 mM ATP, 1 mM ADP, 1 mM FAD, 1 mM NAD+ and either one of the following substrates at a final concentration of 10 mM: [U-^13^C]pyruvate, [U-^13^C]citrate, [U-^13^C]glutamine or [U-^13^C]succinate (Cambridge Isotopes, Inc). For indicated experiments, fluorocitrate (Merck, F9634) was added to a final concentration of 0.5 μM. The nuclei were incubated for one or five hours in the dark at 37°C. Following incubation, the nuclei were centrifuged at 1000g for 8 minutes at 4°C, washed with 1 mL ice-cold PBS and centrifuged again. This step was repeated two times. Polar metabolites were extracted from the nuclei pellet with the addition of 300 μL ice-cold methanol (ULC/MS grade, Biosolve, 136841) supplemented with 10 μL adonitol (50 μm/mL, Alfa AesarTM, L03253.06) as an internal standard and incubation for 15 minutes at 72°C. The methanol/nuclei suspension was further mixed with 300 μL ice-cold MilliQ H2O and centrifuged at 15000 rpm at 4°C for 10 minutes. The supernatants were transferred in amber glass vials (Agilent, 5183-2073), dried with the Genevac EZ-2 Plus evaporator (program, hplc fraction; temperature, 30°C), and stored at −80°C until analysis with Gas Chromatography - Mass Spectrometry.

For the 13C-experiments in whole cell lysates, we followed the same procedure described for the isolated nuclei with slight modifications. In brief, fresh whole cell lysates resuspended in buffer B were adjusted to the incubation buffer by adding the missing compounds and were incubated as above. For the extraction of polar metabolites, 300 μL of whole cell lysates in incubation buffer were mixed with 600 μL ice-cold methanol supplemented with 10 μL adonitol (50 μm/mL) and were incubated for 15 minutes at 72°C. Following the addition of 600 μL ice-cold MilliQ H2O, the rest of the steps were performed as indicated above.

### Gas Chromatography - Mass Spectrometry (GC-MS) data acquisition and analysis

Dried polar metabolites were derivatized with 40 μL of 20 mg/mL methoxyamine hydrochloride (Alfa AesarTM, L08415.14) solution in pyridine (Alfa AesarTM, A12005) for 90 minutes at 37°C, followed by reaction with 80 μL *N*-methyl-*N*-(trimethylsilyl)trifluoroacetamide (Alfa AesarTM, A13141) for 10 hours at room temperature, as justified in (*40*). GC-MS analysis was performed using a Shimadzu TQ8040 GC-(triple quadrupole) MS system (Shimadzu Corp.) equipped with a 30 m x 0.25 mm x 0.25 μm ZB-50 capillary column (Phenomenex, 7HG-G004-11). One μL of sample was injected in split mode (split ratio 1:5 for the isolated nuclei; split ratio 1:30 for the whole cell lysates) at 250°C using helium as a carrier gas with a flow rate of 1 mL/minute. GC oven temperature was held at 100°C for 4 minutes followed by an increase to 320°C with a rate of 10°C/minute, and a final constant temperature period at 320°C for 11 minutes. The interface and the ion source were held at 280°C and 230°C, respectively. The detector was operated both in scanning mode recording in the range of 50-600 m/z, as well as in MRM mode for specified metabolites. For peak annotation the GCMSsolution software (Shimadzu Corp.) was utilized. The metabolite identification was based on an in-house database with analytical standards utilized to define the retention time, the mass spectrum and the quantifying ion fragment for each specified metabolite. The ratio of the different mass isotopomers for each metabolite was determined by integrating the area under the curve (AUC) of the quantifying ion fragments followed by correction for the presence of natural abundant isotopes with the Isotope Correction Toolbox (ICT) (*41*) (related to Tables S1, S2 and S6).

### Nuclear Localization Signal (NLS) motifs prediction

To identify potential NLS motifs inside of the protein sequences of interest (related to Fig. 2G) we used a computational screen. For this purpose, we utilized the sophisticated definitions of NLS motif classes of the ELM database (*42*) and extended the Regular expression search pattern in that way that we allowed further matches by relaxing the pattern restrictions outside of the NLS consensus core motif. For the final pattern search, we used a local version of the pattern search algorithm 3of5 (*43*).

### Generation of constructs with the TCA cycle enzymes fused with the biotin ligase and a FLAG tag

For the selected enzymes, a smaller version of the biotin ligase (BioID2) (*35*) and a FLAG epitope were fused at the C-terminus of the enzymes. For IDH2, we additionally created a version carrying the biotin ligase and the FLAG tag at the N-terminus of the enzyme. Constructs containing the gene of interest fused to the biotin ligase and a FLAG epitope were generated via Gateway cloning technology (Thermo Fisher Scientific). Firstly, we created two destination vectors containing the BioID2 and FLAG for N-terminus fusion and C-terminus fusion. To achieve this, we utilized two destination vectors with the older version of BioID (*44*) (pcDNA5-pDEST-BioID-FLAG-N-term and pcDNA5-pDEST-BioID-FLAG-C-term, kindly donated from Martin Beck (*45*), EMBL Heidelberg) and replaced it with the new and smaller version of BioID2 via Gibson assembly (New England Biolabs). Briefly, BioID2 in pcDNA3.1-BioID2-HA plasmid (Addgene, #74224) was PCR amplified with primers containing the FLAG epitope for N- or C-terminus integration (namely, FLAG-BioID2 or BioID2-FLAG). pcDNA5-pDEST-BioID-FLAG-N-term and pcDNA5-pDEST-BioID-FLAG-C-term plasmids were amplified by PCR, excluding the BioID-FLAG region, and creating appropriate overlapping to the BioID2-FLAG or FLAG-BioID2 end fragments. One μL of the PCR products (15 ng BioID2-FLAG or 15 ng FLAG-BioID2, 7.5 ng pcDNA5-pDEST---N-term or 10 ng pcDNA5-pDEST---C-term) were incubated with 2 μL Gibson assembly reaction mix at 50°C for 60 minutes. Two μL of the reaction were diluted with 8 μL water, and 4 μL of this dilution were used to transform One ShotTM ccd B Survival 2 T1R competent cells (Thermo Fisher Scientific, A10460). Transformants were selected with 33 μg/mL chloramphenicol (Merck, 220551) and 100 μg/mL ampicillin (Merck, 171255). The entry clones with the genes of interests (PDHB, ACO2, IDH2, IDH3G, IDH1 and OGDH) were purchased from the Human ORFeome Collection (Dharmacon). The final constructs were created by performing the LR recombination reaction between the destination vectors and the entry clones according to manufacturer’s instructions. In brief, 1 μL of entry clone (∼100 ng) was combined with 1 μL of destination vector (∼150 ng) and 2 μL of LR Clonase II reaction buffer into a final volume of 10 μL with TE buffer. The reaction was incubated at 25°C for 90 minutes followed by addition of 1 μL proteinase K (Merck, 3115887001) and incubation at 37°C for 10 minutes. One μL of the LR reaction was used to transform DH5α competent E. coli (Thermo Fisher Scientific, EC0112) and transformants were selected with the appropriate antibiotics. All constructs were extracted using the QIAprep Spin Miniprep Kit (Qiagen). The final constructs were sequenced-verified by Sanger sequencing at Eurofins Genomics. All PCR conditions were defined using the http://webpcr.appspot.com/. All vectors, selection antibiotics and primers are listed in Tables S7, S8 and S9.

### Generation of stable inducible Flp-In T-REx HeLa cell lines

Flp-In T-REx *HeLa* cells (*46*) were maintained under standard cultivation conditions (37°C, 5% CO2) in Dulbecco’s modified Eagle medium (DMEM, high glucose, GlutaMAX, Thermo Fisher Scientific, 61965026) supplemented with 10% heat-inactivated fetal bovine serum (FBS, Thermo Fisher Scientific, 26140079) and the addition of zeocin (50 μg/mL, Thermo Fisher Scientific, R25005) and blasticidin (5 μg/mL, Thermo Fisher Scientific, R21001).

The constructs containing the TCA cycle enzymes fused to the biotin ligase and a FLAG tag were stably integrated in the Flp-In T-REx *HeLa* cells using the X-tremeGENE 9 DNA Transfection Reagent (Merck) following the manufacturer’s instructions. Briefly, Flp-In T-REx *HeLa* cells were seeded in 6-well plates (1.8 x 10^4^ cells/well) in cultivation medium without antibiotics. After 24h, the transfection mixture was prepared (consisted of 100 μL Opti-MEM (Thermo Fisher Scientific, 31985062), 3 μL of X-tremeGENE 9 DNA Transfection Reagent, 100 ng of plasmid with the construct of interest and 900 ng of pOG44 (Thermo Fisher Scientific, V600520)), incubated for 15 minutes at room temperature, and added in a drop-wise manner to the cells under gentle shaking. On the third day, the cells were expanded via trypsinization (Trypsin-EDTA, Thermo Fisher Scientific, 25300096) into 150 mm plates, and the following day the medium was changed into cultivation medium with the addition of the selection antibiotics blasticidin (5 μg/mL) and hygromycin (200 μg/mL, Thermo Fisher Scientific, 10687010). The following weeks, fresh cultivation medium with the selection antibiotics was added thrice per week. When large colonies were formed, the cells were trypsinized into the same plates and when 80% confluency was reached they were prepared for cryopreservation.

The parental and stable Flp-In T-REx *HeLa* cell lines were routinely monitored for mycoplasma contamination. The parental Flp-In T-REx HeLa cell line has not been authenticated.

### BioID labeling and cell harvesting

The parental and stable Flp-In T-REx *HeLa* cell lines were seeded in 245mm plates (2.0 x 10^6^ cells/plate) in cultivation medium (described in the previous section) without antibiotics. The cells were allowed to attach for 24 hours and then tetracycline (0.13 μg/mL, Merck, T7660) was added in the medium of the stable cell lines for the induction of protein expression. Following 24 hours, both parental and stable cells were supplemented with biotin (13 μM, Merck, B4639) for protein biotinylation. Finally, after 24 additional hours, the cells were washed with 20 mL PBS, detached with 10 mL trypsin-EDTA and collected with 20 mL cultivation medium. The collected cells were centrifuged at 500 g for 5 minutes at 4°C, re-suspended in 15 mL PBS (4°C) and centrifuged. Once the supernatant was aspirated, the cell pellets were frozen on dry ice and stored at −80°C until further processing. For each case, four biological replicates were collected.

### Affinity purification of biotinylated proteins

The affinity purification of the biotinylated proteins was adapted from the method developed by Mackmull et al 2017 (*45*) with slight modifications. The frozen cell pellets were re-suspended in 8 mL of ice-cold lysis buffer with the following composition: 50 mM Tris pH 7.5, 150 mM NaCl, 1% Triton X-100, 0.1% SDS, 1 mM EDTA, 1mM EGTA, 1 mM PMFS (Merck, P7626), 1 mg/mL aprotinin (Carl-Roth, A162.1), 0.5 mg/mL leupeptin (Carl-Roth, CN33.3), 250 Units HS-Nuclease (MoBiTec, GE-NUC10700-01). Following re-suspension, the samples were incubated for one hour at 4°C under constant mild rotation (30 rpm). Subsequently, the cell lysates were sonicated at 4°C for 30 seconds, 5 times, with 30 seconds rest in-between, and were further centrifuged at 17000g for 30 minutes at 4°C to remove any insoluble material. 80 μL of Streptavidin Sepharose High Performance beads (GE Healthcare, 17-5113-01) were equilibrated in 1 mL of lysis buffer for 30 minutes at 4°C under constant mild rotation (30 rpm). The equilibrated beads were centrifuged at 2000g for 5 minutes at 4°C, then transferred to the lysed supernatants and incubated for 3 hours at 4°C under constant mild rotation (30 rpm). After the incubation, the beads were centrifuged at 2000g for 5 minutes at 4°C, and finally 7.5 mL of the supernatant were discarded. The remaining beads-lysates were transferred to a Spin Column (PierceTM, Thermo Fisher Scientific, 69705), washed once with 800 μL of lysis buffer, and then five times with 700 μL of 50 mM ammonium bicarbonate pH 8.3. After the washing, the column was plugged and the beads were transferred to a fresh eppendorf tube with 300 μL of 50 mM ammonium bicarbonate pH 8.3. The same procedure was repeated two more times to ensure all beads were collected. One μg of Sequencing Modified Trypsin (Promega, V5117) was added and the samples were incubated at 37 °C for 16 hours under constant shaking (500 rpm). The following day, 0.5 μg of trypsin were added and the beads were incubated for 2 additional hours under the same conditions. Following incubation, the beads were transferred to a Spin Column and the digested peptides were eluted with 150 μL of 50 mM ammonium bicarbonate pH8. This step was performed one additional time. The eluted peptides were dried in a speed-vac and stored at −80°C until further processing. All steps were performed using low retention tips (TipOne® RPT Tips, Starlab) and low protein binding collection tubes (Thermo Fisher Scientific, 90411).

### Sample preparation for mass spectrometry

Dried samples were dissolved in 1% formic acid with 4% acetonitrile and subjected to OASIS® HLB μElution Plate (Waters) for desalting according to manufacturer’s instructions. Desalted peptides were reconstituted in 50 mM HEPES (pH 8.5) and labelled with TMT10plex158 Isobaric Label Reagent (Thermo Fisher Scientific) according the manufacturer’s instructions. For further sample clean up an OASIS® HLB μElution Plate (Waters) was used. Offline high pH reverse phase fractionation was carried out on an Agilent 1200 Infinity high-performance liquid chromatography system, equipped with a Gemini C18 column (3 μm, 110 Å, 100 x 1.0 mm, Phenomenex) (*47*).

### Mass spectrometry data acquisition

An UltiMate 3000 RSLC nano LC system (Dionex) fitted with a trapping cartridge (μ-Precolumn C18 PepMap 100, 5μm, 300 μm i.d. x 5 mm, 100 Å) and an analytical column (nanoEase™ M/Z HSS T3 column 75 μm x 250 mm C18, 1.8 μm, 100 Å, Waters). Trapping was carried out with a constant flow of solvent A (0.1% formic acid in water) at 30 μL/minute onto the trapping column for 6 minutes. Subsequently, peptides were eluted via the analytical column with a constant flow of 0.3 μL/minute with increasing percentage of solvent B (0.1% formic acid in acetonitrile) from 2% to 4% in 4 minutes, from 4% to 8% in 2 minutes, then 8% to 28% for a further 96 minutes, and finally from 28% to 40% in another 10 minutes. The outlet of the analytical column was coupled directly to a QExactive plus (Thermo Fisher Scientific) mass spectrometer using the proxeon nanoflow source in positive ion mode.

The peptides were introduced into the QExactive plus via a Pico-Tip Emitter 360 μm OD x 20 μm ID; 10 μm tip (New Objective) and an applied spray voltage of 2.3 kV. The capillary temperature was set at 320°C. Full mass scan was acquired with mass range 350-1400 m/z in profile mode in the FT with resolution of 70000. The filling time was set at maximum of 100 ms with a limitation of 3×10^6^ ions. Data dependent acquisition (DDA) was performed with the resolution of the Orbitrap set to 35000, with a fill time of 120 ms and a limitation of 2×10^5^ ions. A normalized collision energy of 32 was applied. A loop count of 10 with count 1 was used and a minimum AGC trigger of 2e2 was set. Dynamic exclusion time of 30 s was used. The peptide match algorithm was set to ‘preferred’ and charge exclusion ‘unassigned’, charge states 1, 5 - 8 were excluded. MS2 data were acquired in profile mode (*48*).

### Mass spectrometry data analysis

IsobarQuant (*49*) and Mascot (v2.2.07) were used to process the acquired data, which was searched against a UniProt Homo sapiens proteome database (UP000005640) containing common contaminants and reversed sequences. The following modifications were included into the search parameters: Carbamidomethyl (C) and TMT10 (K) (fixed modification), Acetyl (N-term), Oxidation (M) and TMT10 (N-term) (variable modifications). For the full scan (MS1) a mass error tolerance of 10 ppm and for MS/MS (MS2) spectra of 0.02 Da was set. Further parameters were set: Trypsin as protease with an allowance of maximum two missed cleavages: a minimum peptide length of seven amino acids; at least two unique peptides were required for a protein identification. The false discovery rate on peptide and protein level was set to 0.01.

### Identification of putative interactors for each bait enzyme of TCA cycle

Raw data of IsobarQuant were loaded into R (ISBN 3-900051-07-0). As a quality criteria, only proteins which were quantified with at least two different unique peptides were used for downstream analysis. The “signal_sum” columns of the “proteins”-output sheet from IsobarQuant were cleaned for potential batch-effects with limma (*50*) and subsequently normalized with vsn (variance stabilization) (*51*). Missing values were imputed with the impute function (method = “knn”) from the MSNBase package (*52*). To create the overview of the biotinylated proteins significantly associated with each bait enzyme, hereafter termed as overview of the putative interactors (related to Fig. 3A and Table S3), the following two comparisons were made. Firstly, each cell line expressing a bait enzyme with the biotin ligase fused at the C-terminus was compared to parental cells using limma. A biotinylated protein was considered a putative interactor with a false discovery rate (fdr) ≤ 20 % and a fold change of at least 50 %. These results were subsequently refined by removing proteins characterized as contaminants in the CRAPome database (*53*) (version 1.1) with the following parameters: organism, H. sapiens; cell/tissue type, *HeLa*; epitope tag, BirA*-FLAG; cutoff frequency, detected in more than 6 experiments (out of 16 experiments in total), unless they displayed a fold change (engineered Flp-In T-REx *HeLa* to parental Flp-In T-REx *HeLa*) ≥ 2.5. Secondly, each cell line expressing a bait enzyme fused with biotin ligase at the C-terminus was compared to the cell line expressing IDH2 carrying the biotin ligase at the N-terminus. This fusion renders the mitochondrial import signal at the N-terminus of IDH2 inaccessible; as such, this engineered enzyme cannot access the mitochondria, rather resides in the cytoplasm (related to Table S10), and serves as a supplemental negative control to further eliminate unspecific biotinylated proteins as a result of the exogenous expression of the biotin ligase. A biotinylated protein was considered a putative interactor using the same analysis and criteria as in the first comparison. Finally, the overview included only those defined as putative interactors in both comparisons.

To define for each bait enzyme a subset of putative interactors that show significant enrichment as compared to the mitochondrial-only IDH2 enzyme (related to Fig. 3B and Table S5), hereafter termed as top putative interactors, we next compared each cell line expressing a bait enzyme with the biotin ligase fused at the C-terminus to: a) the parental cells, b) the cells expressing IDH2 with the biotin ligase fused at the N-terminus, and c) the cells expressing the IDH2 carrying the biotin ligase at the C-terminus. In each of the three comparisons, a biotinylated protein is considered a putative interactor using the same analysis and criteria as in the above paragraph. Finally, for each bait enzyme, the proteins defined as putative interacting proteins in all three comparisons were retained. For the case of IDH2 with the biotin ligase at the C-terminus, the three following comparisons were performed: a) to the parental cells, b) to the cells expressing the IDH2 with the biotin ligase fused at the N-terminus, and c) to the cells expressing ACO2 with the biotin ligase fused at the C-terminus.

### Heat map and clustering analysis

Clustering analysis (related to Fig. 3A) was performed for the overview of the putative interactors associated with each bait enzyme (Table S3). The input values for each putative interactor corresponded to the average (across all biological replicates) log2-fold changes in the abundance of the putative interactor in cells expressing a bait enzyme relative to the parental *HeLa* cells. Hierarchical clustering was performed on the centered and scaled input values with the “eclust” package (*54*) using the “ward.D2” linkage and the “Euclidean” distance metric. For visualization, the “ComplexHeatmap” package (*55*) was utilized in R.

### Gene ontology enrichment analysis

The five major clusters of the putative interactors defined by the hierarchical clustering (related to Fig. 3A), were subjected to gene ontology enrichment analysis (related to Tables S4 and S10) by employing g:Profiler (*56*) (version 0.1.9). The following parameters were defined: organism, “hsapiens”; significance threshold correction method, “g:SCS”; user threshold, “0.05”. The statistical domain scope was set to “custom” and the entire set of detected putative interacting proteins was used as a background list. For the case of IDH2 carrying the biotin ligase at the N-terminus (related to Table S10), gene ontology enrichment analysis was performed using the “annotated” genes for the statistical domain scope since the “custom” list did not result in statistically significant results.

### Topological distribution of the biotinylated proteins

To define the topological distribution (related to Fig. 3B) of the top putative interactors for each of the bait enzymes (Table S5), we retrieved information from UniProt database (*57*) (downloaded on 14/09/2020) and the Human Protein Atlas (*31*) (version 19.3, available from http://www.proteinatlas.org). UniProt was queried for: i) the subcellular localization manual assignments, and ii) for the Gene Ontology annotations for the term “Cellular Component”. The input data corresponded to the respective reviewed entries for H. sapiens. Human Protein Atlas was queried on the subcellular location data.

### Dotplot generation

For the dotplot generation (related to Fig. 3E), the ProHits-viz platform (*58*) was used for the top putative interactors (Table S5). The input data corresponded to: i) the average (across all biological replicates) log2-fold changes in the abundance of each biotinylated protein in the cells expressing a bait enzyme relative to the parental *HeLa* cells, and ii) the respective p values estimated with limma analysis. The following parameters were used: score type, p value defined by limma analysis; primary filter, 0.01; secondary filter, 0.1; no clustering was utilized.

### Immunofluorescence labeling

For the immunofluorescence analysis of the engineered Flp-In T-REx *HeLa* expressing a bait enzyme (related to Fig. 3, C and D, Fig. 4A, fig. S3, fig. S4 and fig. S5), cells were seeded in coverglass chambers (Nunc Lab-Tek II, Merck, Z734853) and allowed to attach for 24 hours prior to tetracycline addition (0.13 μg/mL, Merck, T7660) for the induction of protein expression. After 24 hours, biotin (13 μM, Merck, B4639) was added for protein biotinylation. Finally, after 24 additional hours, the cells were washed with PBS and fixed for 10 minutes with 4% formaldehyde (Image-iT fixative solution, Thermo Fisher Scientific, FB002) at room temperature. Following three washing steps with PBS, the cells were permeabilized for 20 minutes with 0.2% Triton X-100 in PBS, washed thrice with PBS, and blocked (0.1% BSA, 0.3M glycine, 0.1% Tween 20, in PBS) for 1h at room temperature. Incubation with primary antibodies (0.3M glycine, 0.1% Tween 20, in PBS) was performed overnight in a humidified dark chamber at 4°C. The following antibodies and dilutions were used: streptavidin conjugated to Alexa Fluor 488 (1:100, Thermo Fisher Scientific, S11223), rabbit anti-OGDH (1:100, Merck, HPA020347), rabbit anti-IDH3G (1:100, Merck, HPA002017), rabbit anti-ACO2 (1:100, Abcam, ab129069), rabbit anti-IDH2 (1:100, Merck, HPA007831), mouse anti-Tom20 (1:100, BD Biosciences, 612278). The next day, the cells were washed three times with PBS and 0.1% (v/v) Tween 20 (PBST), and incubated with secondary antibodies (0.3M glycine, 0.1% Tween 20, in PBS) for 1 hour in a humidified dark chamber at room temperature. The following secondary antibodies and dilutions were used: anti-rabbit Alexa Fluor 647 (1:200, Thermo Fisher Scientific, A27040) and anti-mouse Alexa Fluor 555 (1:200, Thermo Fisher Scientific, A28180). Following three washing steps with PBST, the cells were stained with Hoechst (2.6 μg/mL, Thermo Fisher Scientific, 62249) for 20 minutes in the dark at room temperature. Finally, the cells were washed three times with PBS and saved in PBS supplemented with 0.02% sodium azide (Merck, S2002) at 4°C until image acquisition.

For the immunofluorescence analysis of cell lysates and isolated nuclei (related to Fig. 4B and fig. S6), engineered Flp-In T-REx *HeLa* cells were grown and treated with tetracycline and biotin as before, followed by the nuclei isolation procedure described above. Cell lysates and isolated nuclei were resuspended in buffer (0.25 M sucrose, 50 mM Tris-HCl pH 7.5, 25 mM KCl, 5 mM MgCl2, 2 mM DTT, 10 mg/mL leupeptin, 5 mg/mL aprotinin) and were attached via centrifugation on coverslips (thickness #1.5, Thermo Fisher Scientific, CB00110RAC20MNT0) previously coated with poly-L-lysine (Thermo Fisher Scientific, P4832). The rest of the steps were the same as described for the immunofluorescence analysis of whole cells with the following modifications: primary and secondary antibodies incubation buffer consisted of 0.3M glycine and 0.025% Tween 20 in PBS, Hoechst staining was performed for 10 minutes, and all washes were done with PBS. Finally, all coverslips were mounted on slides with one drop of ProLong Gold Antifade mountant (Thermo Fisher Scientific, P10144), dried in the dark overnight at room temperature, and saved at -20°C until image acquisition. The following primary and secondary antibodies along with the respective dilutions were used: streptavidin conjugated to Alexa Fluor 488 (1:100, Thermo Fisher Scientific, S11223), mouse anti-lamin B1 (1:500, Atlas Antibodies, AMAb91251) and anti-mouse Alexa Fluor 555 (1:1000, Thermo Fisher Scientific, A28180).

For the immunofluorescence analysis of mouse embryonic stem (ES) cells (related to Fig. 4, C and D, and fig. S7), cells were grown on mouse embryonic fibroblasts (MEFs) in KnockOut™ DMEM medium (Gibco, 10829018) supplemented with 1% Penicillin-Streptomycin (Gibco, 15070063), 1% L-glutamine (Gibco, 25030081), 1% MEM Non-Essential Amino Acids Solution (Gibco, 11140050), 15% FBS (Gibco, 10270106), 0.024 μg/mL of LIF (EMBL Protein expression and purification core facility) and 0.12mM of 2-Mercaptoethanol (Gibco). Cells were maintained on MEFs and incubated at 37°C with 5% CO2 and 95% relative humidity. TrypLE™ Express Enzyme (1X) (Gibco, 12605036) was used to detach cells for passaging, collection or to create a single cell suspension. For immunofluorescence staining, 31.600 ES cells per well were grown without feeders on chambers (ibidi, µ-slide 8 well, #80826) previously coated with 0.1% gelatin in 300µl of growth medium. After 48 hours, ES cells were subjected to immunofluorescence staining following the same procedure as for the engineered Flp-In T-REx *HeLa* cells.

### Image acquisition and analysis

Images of the engineered Flp-In T-REx *HeLa* cells (related to Fig. 3, C and D, Fig. 4A, fig. S3, fig. S4 and fig. S5) were acquired as single optical sections using the same settings for all conditions at the Advanced Light Microscopy Facility at EMBL, Heidelberg, with a Leica TCS SP8 system and a 63x/1.40 oil objective (HC PL APO CS2). To capture the fluorophores and eliminate any cross talk, the following three sequential acquisition settings with the respective wavelength excitation (ex) and emission (em) detection windows were utilized: Nr.1, for Hoechst and Alexa Fluor 647, ex:405/em:420-490 nm and ex:638/em:650-750 nm, respectively; Nr.2, for Alexa Fluor 488, ex:488/em:500-531 nm; Nr.3, for Alexa Fluor 555, ex:552/em:562-592 nm. Scan speed was set at 100 Hz, line averaging at 5, and image resolution was 3144×3144 pixels. The image files were subsequently processed with ImageJ/Fiji (National Institutes of Health, NIH) software (*59*). Briefly, nuclei were located based on the Hoechst staining and a region-of-interest (ROI) inside each nucleus was manually selected to ensure that only pixels inside the nucleoplasm are measured. Next, a ROI covering the nucleus and an extended area around was manually created delineating the “whole cell”. Finally, staining based on Tom20, an outer mitochondrial membrane protein, was utilized to segment the “mitochondria” ROIs. The biotinylation levels were quantified in whole cells (related to Fig. 3D and fig. S3B), mitochondria regions (related to fig. S3C) and the nuclei regions (related to Fig. 4A) based on the mean intensity values of Alexa Fluor 488-conjugated streptavidin staining. To quantify the enzyme levels, the mean intensity of the enzyme was measured in the “whole cell” regions (related to Fig. 3C and fig. S3A). The mean intensity values were examined for statistical significance with the Wilcoxon rank sum test. For visualization purposes, all images were treated using the same intensity range (min/max) (related to fig. S4 and S5).

Images of lysed cells and isolated nuclei from the engineered Flp-In T-REx *HeLa* cells (related to Fig. 4B and fig. S6) were acquired as described above with the modification that Nr.1 acquisition setting included only the Hoechst wavelength excitation and emission detection window and image analysis was performed only for the “nucleus” ROIs. To account for possible technical variations, we further normalized the biotinylation levels in each individual ROI to the corresponding mean intensity values of the Hoechst staining. Statistical analysis and visualization were performed as described in the section above (related to Fig. 4B and fig. S6).

Images of mouse embryonic stem cells (related to Fig. 4, C and D, and fig. S7) were acquired in the form of z-stacks (step size of 0.34 µm) using the same settings for all conditions with a Confocal Leica TCS SP5 microscope and a 40.0×1.25 oil objective (HC PL Apo UV optimized). The following two sequential acquisition settings with the respective wavelength excitation (ex) and emission (em) detection windows were utilized: Nr.1, for Hoechst, ex:405/em:420-612 nm; Nr.2, for Alexa Fluor 647, ex:638/em:644-800 nm. Scan speed was set at 100 Hz and image resolution was 1024×1024 pixels. The three-dimensional image reconstruction and downstream analysis were performed with the Imaris v.9.5.1 (Bitplane) software. In brief, the “surfaces” function was utilized with the Hoechst staining as the source channel to create 3D surfaces demarcating the nuclei. To ensure that the regions covered mainly the nucleoplasm, a high threshold value (absolute threshold option set at 100; smoothing option disabled) was selected. Subsequently, the mean intensity values of the fluorescent channel corresponding to the enzymes staining (Alexa Fluor 647) were measured within the 3D nuclei surfaces (related to Fig. 4, C and D, and fig. S7, A and C). To quantify the enzyme levels in the mitochondria (related to fig. S7B), we utilized the “spots” function for the enzyme staining channel. The different spot sizes option was selected, the diameter was set at 1 µm, and background subtraction was enabled. To capture the spots relevant to the mitochondria regions, we used the mean intensity of the source channel as the filter type with a high threshold value (50). Finally, the sensitivity for defining the spots was further adjusted using the “spots region from local contrast” option and a high threshold value (100). The mean intensity values were examined for statistical significance with the Wilcoxon rank sum test. For visualization purposes, one representative plane is depicted while all images were treated using the same intensity range (min/max) (related to Fig. 4D and fig. S7A).

### Immunoblotting

For immunoblotting (related to Fig. 1B, fig. S8, fig. S9 and fig. S10), isolated nuclei were collected as described in the respective section above. The samples were lysed in SDS-PAGE sample buffer (62.5 mM Tris pH 6.8, 2% SDS, 10% glycerol, 0.0006% Bromophenol Blue, 5% β-Mercaptoethanol), incubated at 95°C for 5 minutes, and finally sonicated to shear the DNA. The proteins were separated on 4–20% gradient gels (Mini-PROTEAN® TGX™ Precast Protein Gels, Biorad). Prior transfer, the gel, the nitrocellulose membrane (Thermo Fisher Scientific, 88018) and the blot filter papers (Biorad, #1703932) were equilibrated in transfer buffer (25 mM Tris, 192 mM glycine pH 8.3, 20% methanol, 0.04 % SDS) for 15 minutes with agitation. For the transfer, the Trans-Blot Turbo System (Biorad) with the “STANDARD SD” protocol followed by the “Mixed MW” program was used. The membrane was stained with Amido Black solution (Merck, A8181) for 10 minutes and imaged in a ChemiDoc MP Imaging System (Biorad). Following de-staining with water, the membrane was blocked with 3% BSA in PBST for one hour at room temperature. The primary antibodies were diluted in 5% BSA in PBST and the membrane was incubated overnight at 4°C under agitation. The following day, the membrane was washed thrice with PBST, incubated for one hour at room temperature with the appropriate secondary antibodies in 3% BSA in PBST, and then washed thrice with PBST. Finally, the membrane was developed using the Pierce ECL Plus Western Blotting Substrate (Thermo Fisher Scientific, 32132) and imaged in a ChemiDoc MP Imaging System (Biorad). The following antibodies and dilutions were used: rabbit anti-IDH2 (1:1000, Merck, HPA007831), rabbit anti-COXIV (1:1000, Cell Signaling Technology, #8674), rabbit anti-cytochrome c (1:1000, Cell Signaling Technology, #8674), rabbit anti-beta III tubulin (1:1000, Abcam, ab18207) and anti-rabbit HRP-linked (1:2000, Abcam, ab205718). As a molecular weight marker we utilized the Precision Plus Protein™ Prestained Standard in Dual Color (Biorad, 1610374).

## Supplementary Figures

**Fig. S1.**
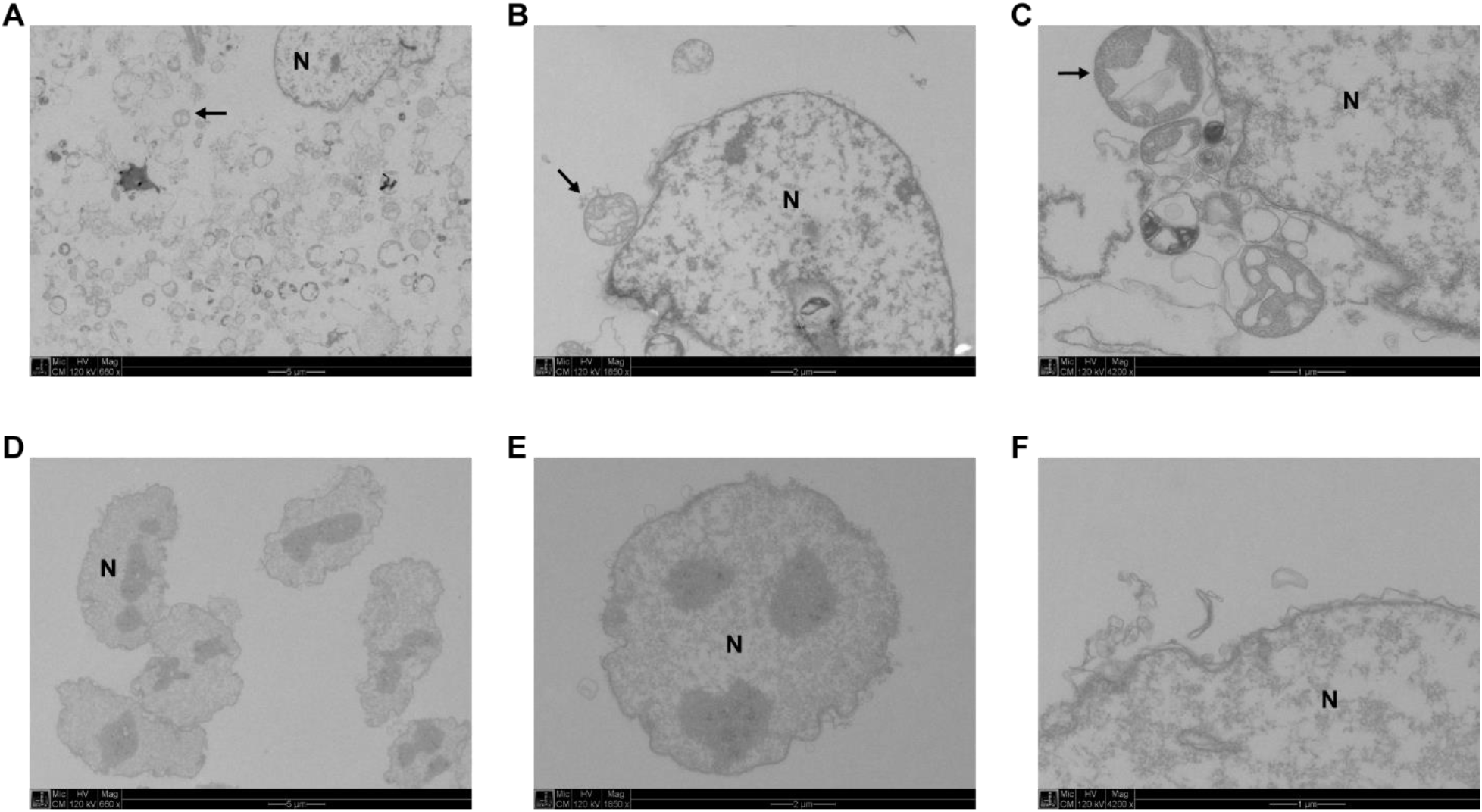
Representative electron microscopy images from lysed cells (A to C) and isolated nuclei (D to F). For both lysed cells and isolated nuclei, the following magnifications are presented: x660 (A and D), x1850 (B and E) and x4200 (C and F). The letter “N” indicates a representative nucleus. The arrows point to mitochondria.

**Fig. S2.**
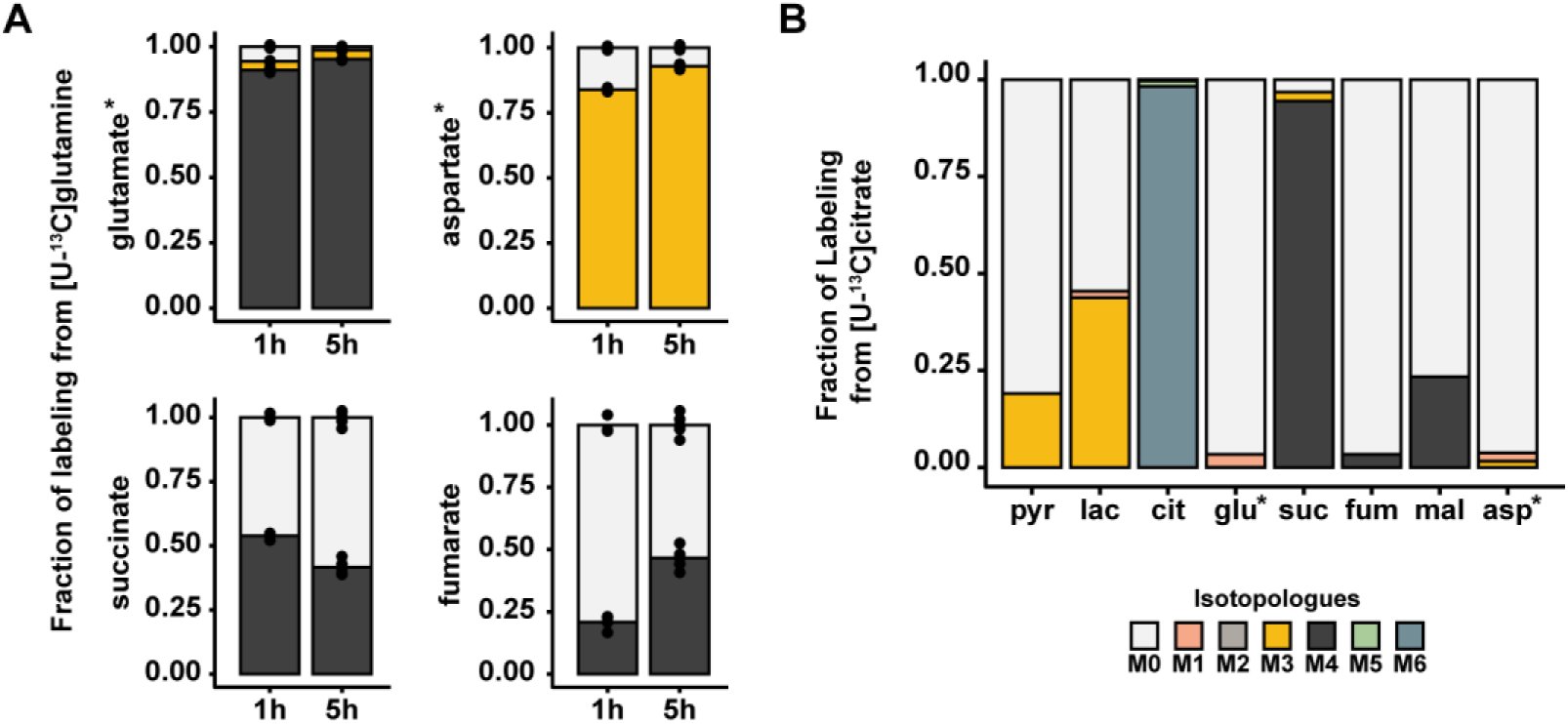
(A) Fraction of labeling of the different mass isotopologues for glutamate, aspartate, succinate and fumarate in nuclei incubated for 1 (1h) and 5 hours (5h) with [U-13C]glutamine. For the mass isotopologues (Mn; n, number of 13C-carbons) legend refer to panel B. For the 1 hour incubation, n = 3 biological replicates. For the 5 hours incubation, n = 6 biological replicates. (B) Fraction of labeling of the different mass isotopologues (Mn; n, number of 13C-carbons) for each metabolite in lysed cells incubated with [U-13C]citrate for 5 hours. One representative experiment is depicted. In panels A and B, the quantified ions for glutamate and aspartate correspond to a four and a three carbon fragment, respectively. Data are presented as mean of the indicated biological replicates with individual data points shown. Abbreviations: pyr, pyruvate; lac, lactate; cit, citrate; glu, glutamate; suc, succinate; fum, fumarate; mal, malate; asp, aspartate.

**Fig S3.**
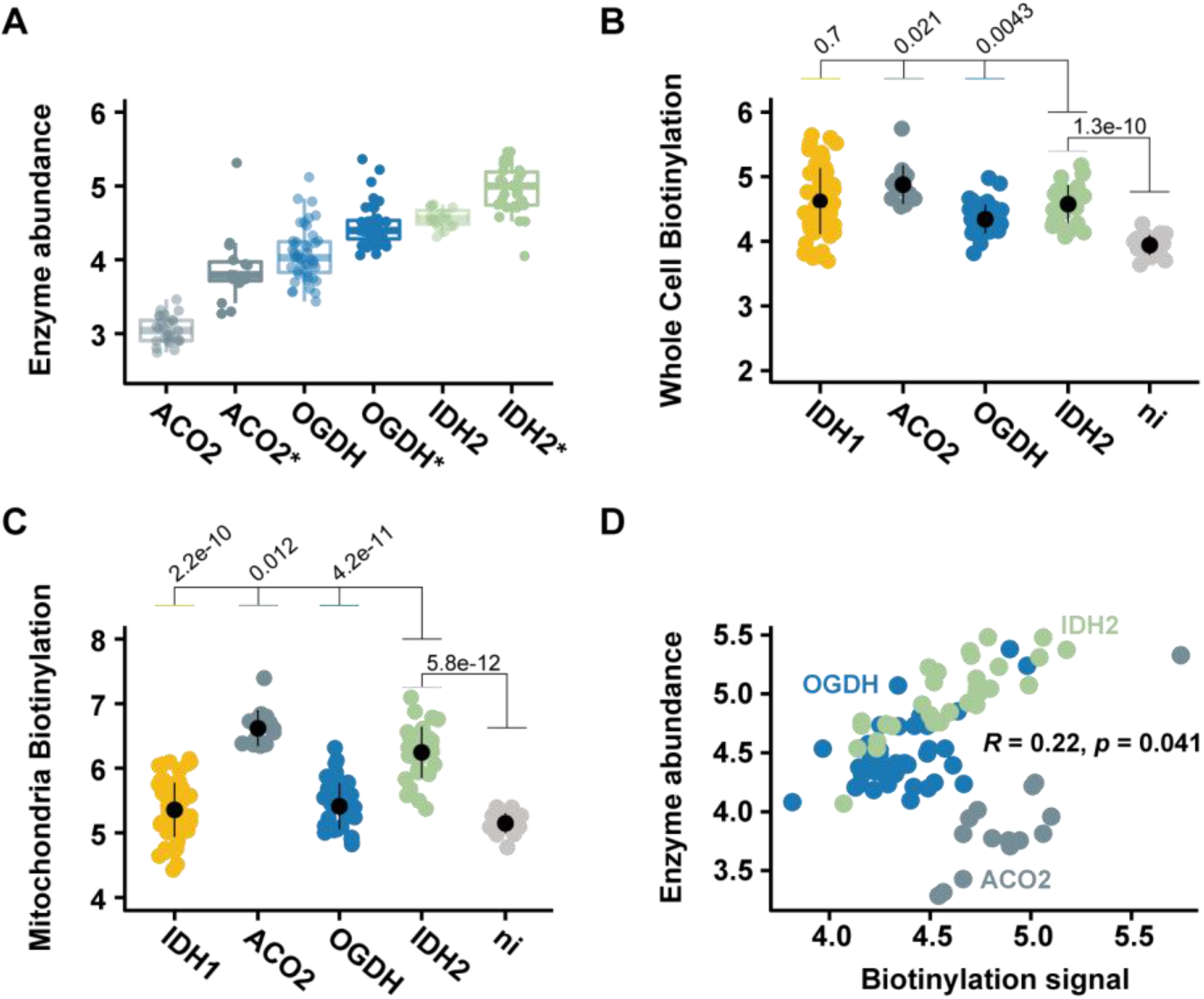
(A) Abundance levels of the bait enzymes before and after (demarcated by *) induction of their expression. The quantification was based on immunofluorescence microscopy using antibodies specific to the respective endogenous enzymes. From left to right, n = 27, 15, 45, 40, 19 and 31 cells quantified per case. (B and C) Quantification of the biotinylated proteins in the whole cell area (B) and in the mitochondria (C) of cells expressing the indicated bait enzymes. The assessment was based on immunofluorescence microscopy using fluorescently-labeled streptavidin. “ni” refers to a cell line where the expression of the bait enzyme and the biotinylation was not induced; for the particular case we used the cell line of OGDH. For panels B and C, from left to right, n = 60, 15, 40, 31 and 19 quantified cells. (D) Pearson correlation analysis of the enzyme abundance levels (defined in panel A) with the biotinylation levels (defined in panel B) in cells expressing the indicated bait enzymes. For ACO2, n = 15; for OGDH, n = 40; for IDH2, n = 31. In panels (A) to (D), points represent quantification (log2 scale) of individual cells with the population mean and standard deviation calculated for each case. In panels (A) to (C), significance was assessed for the indicated comparisons using the Wilcoxon rank sum test.

**Fig S4.**
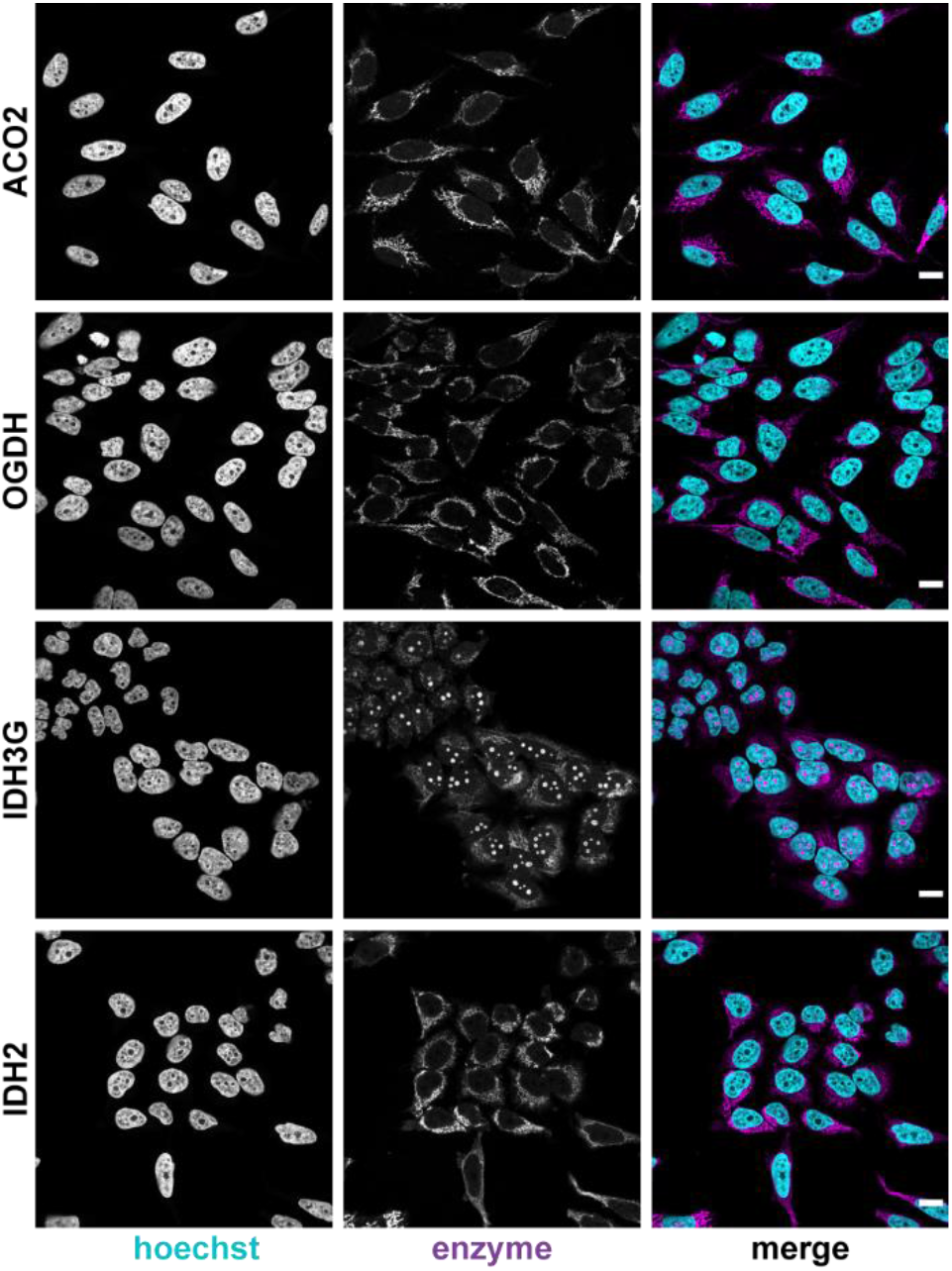
Subcellular distribution of the bait enzymes in HeLa cells. The localization was detected using antibodies specific to the endogenous enzymes (magenta). DNA was stained with Hoechst (cyan). Scale bars, 15µm.

**Fig S5.**
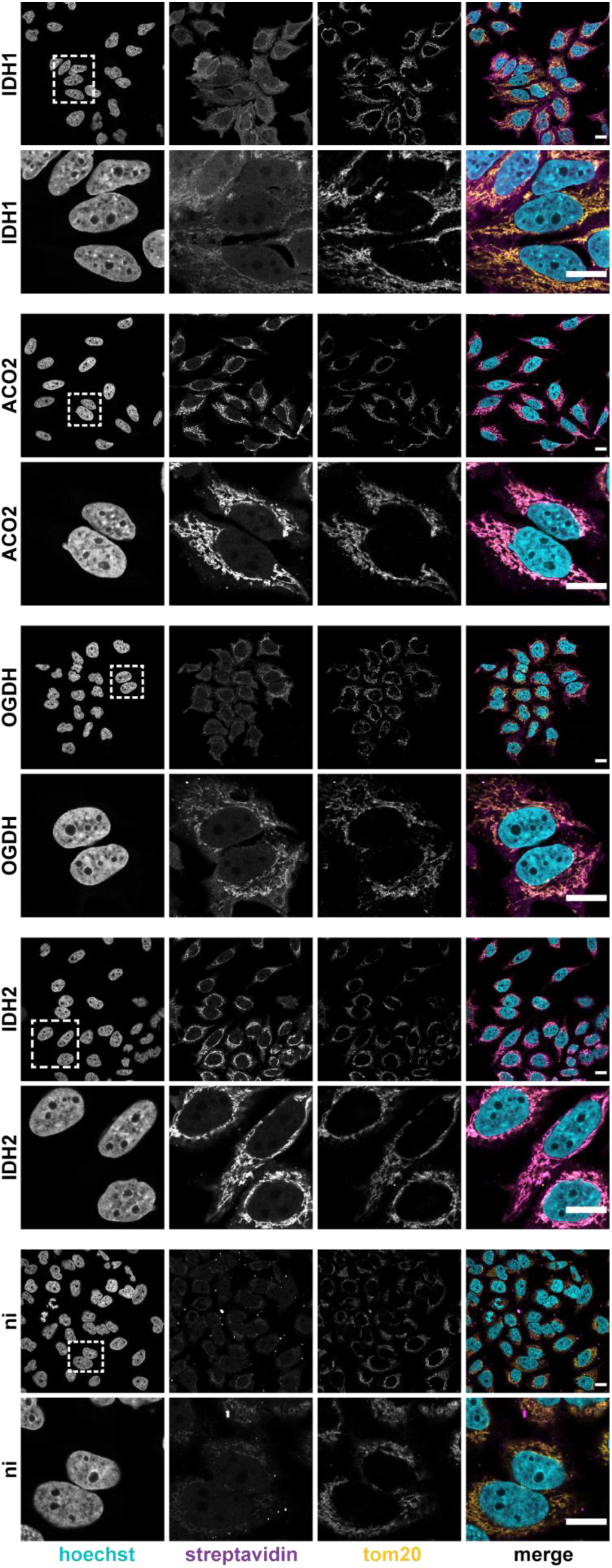
Subcellular distribution of the biotinylated proteins in the cell lines expressing the bait enzymes. The biotinylation was detected using fluorescently-labeled streptavidin (magenta). Cells were also stained for the outer mitochondrial membrane protein Tom20 (yellow), and DNA (cyan). For each case, a wide-field (upper panel) and a close-up (lower panel) view are depicted. “ni” refers to cells where the expression of a bait enzyme and the biotinylation has not been induced. Scale bars, 15µm.

**Fig S6.**
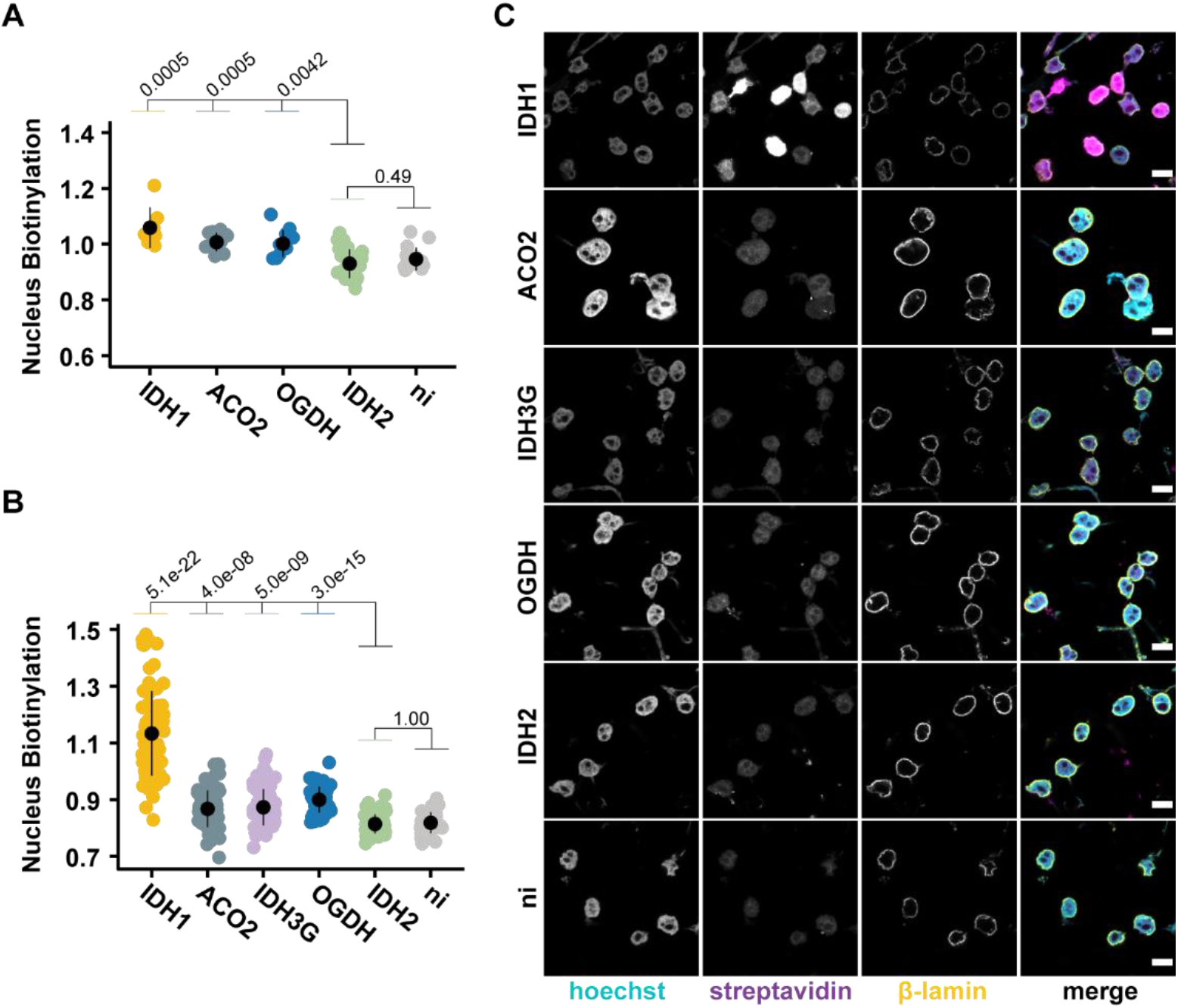
(A and B) Quantification of the biotinylated proteins in the nucleus region of lysed cells (A) and in isolated nuclei (B) from cells expressing the bait enzymes. The quantification was based on immunofluorescence microscopy using fluorescently-labeled streptavidin. “ni” refers to cells where the expression of a bait enzyme and the biotinylation has not been induced. Points represent quantification (log2 scale) of individual nuclei with the population mean and standard deviation calculated for each case. Significance was assessed for the indicated comparisons using the Wilcoxon rank sum test. For panel A, from left to right, n = 7, 12, 11, 24 and 15 quantified nuclei. For panel B, from left to right, n = 73, 75, 69, 55, 63 and 37 quantified nuclei. (C) Representative immunofluorescence images of nuclei isolated from cells expressing the bait enzymes. The nuclei were stained for the biotinylated proteins with fluorescently-labeled streptavidin (magenta), for the nuclear membrane protein β-lamin (yellow) and for DNA (cyan). Scale bars, 15µm.

**Fig S7.**
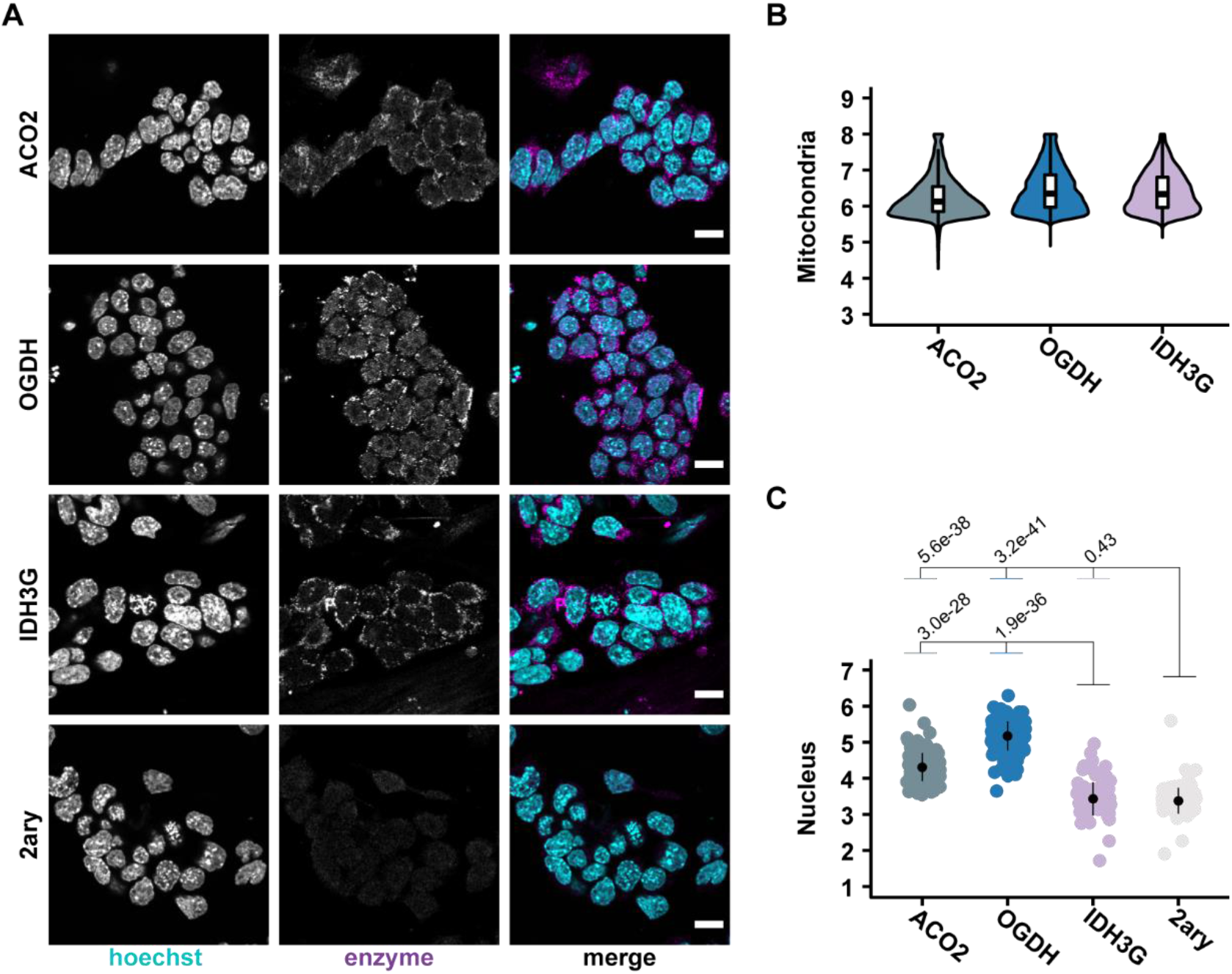
(A) Representative immunofluorescence images of embryonic stem cells stained for ACO2, OGDH and IDH3G. The detection was based on antibodies specific to the enzymes (magenta). DNA was stained with Hoechst (cyan). “2ary” refers to cells stained only with the secondary, fluorophore-conjugated, antibody. One representative plane is depicted. Scale bars, 15µm. (B and C) Quantification of the enzymes levels in the mitochondria (B) and in the nuclei (C) of embryonic stem cells. In (C), points represent quantification (log2 scale) of individual nuclei with the population mean and standard deviation calculated for each case. Significance was assessed for the indicated comparisons using the Wilcoxon rank sum test. For panel B, from left to right, n = 7467, 18457 and 2846 quantified mitochondria spots. For panel C, from left to right, n = 183, 166, 132 and 133 quantified nuclei.

**Fig S8.**
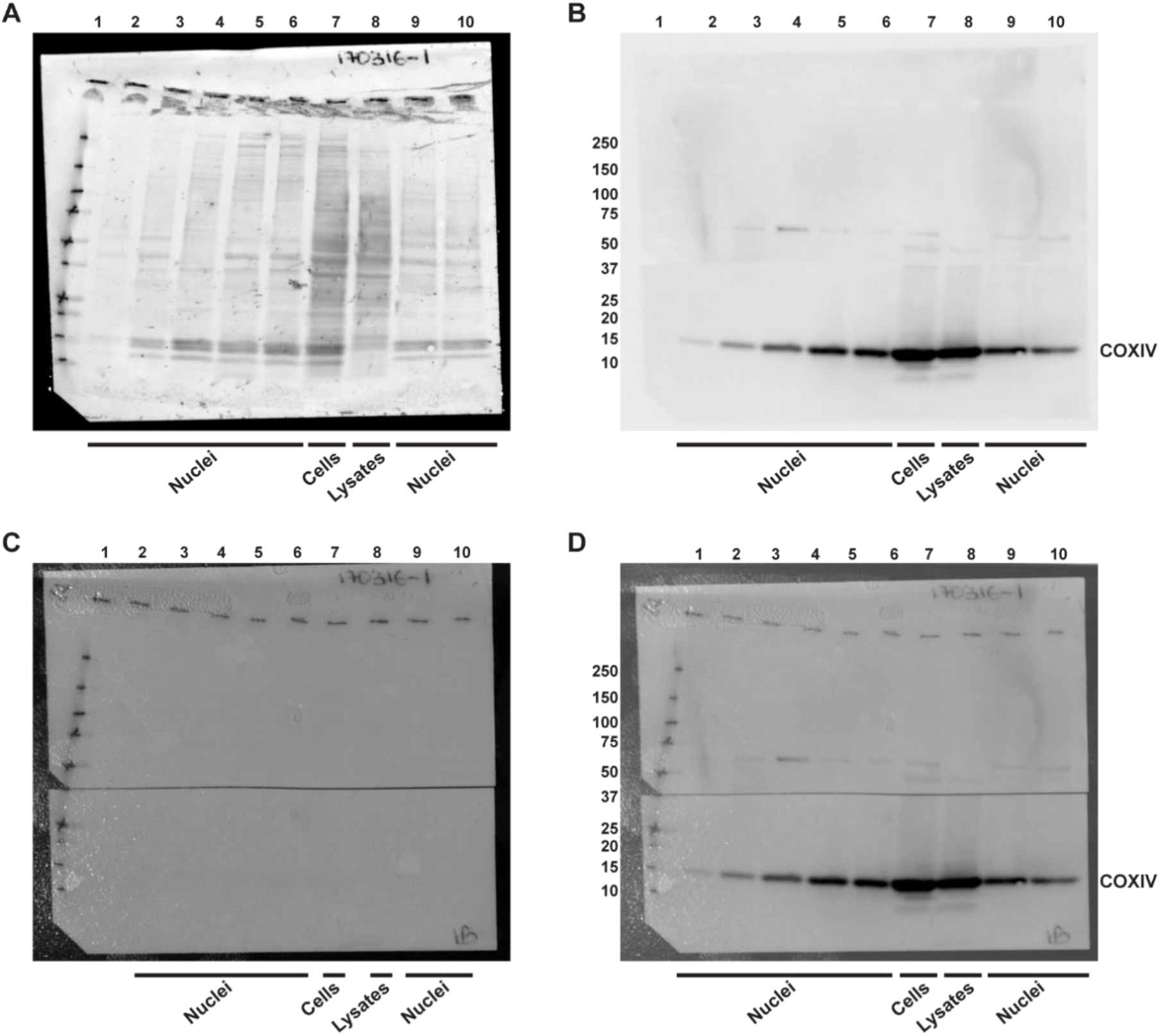
Immunoblot analysis for the inner mitochondrial membrane protein, COXIV, in isolated nuclei, cells and whole cell lysates (related to Fig. 1B). (A) Amido black staining of the membrane for detection of total proteins. (B) The membrane was cut in two pieces and the lower part was incubated with the antibody specific to COXIV. The chemiluminescent image of the membrane is displayed. The numbers on the left side of the panel correspond to the molecular weight (kD) of the protein standards. (C) Colorimetric image of the membrane. (D) Overlay of the chemiluminescent (panel B) and colorimetric (panel C) images. Well 1, protein standards (10 to 250 kD); wells 2 to 5, isolated nuclei; well 6, cells; well 7, cell lysates; wells 8 to 9, isolated nuclei. The wells loaded with nuclei samples correspond to independent isolations.

**Fig S9.**
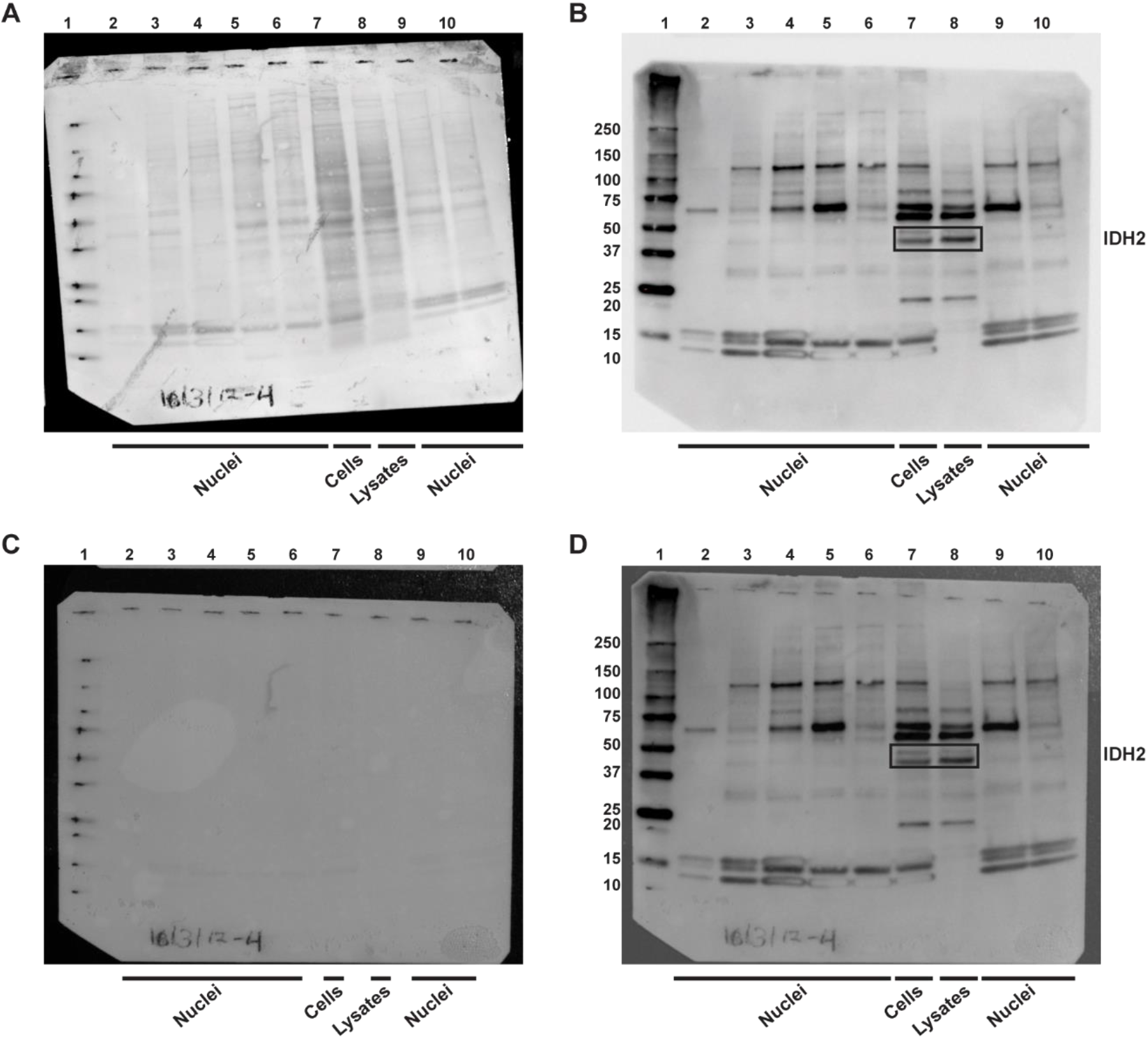
Immunoblot analysis for the mitochondrial matrix protein, IDH2, in isolated nuclei, cells and whole cell lysates (related to Fig. 1B). (A) Amido black staining of the membrane for detection of total proteins. (B) Chemiluminescent image of the membrane following incubation with antibody specific to IDH2. The band that correspond to IDH2 according to the expected molecular weight of the enzyme are highlighted with a black box. The numbers on the left side of the panel correspond to the molecular weight (kD) of the protein standards. (C) Colorimetric image of the membrane. (D) Overlay of the chemiluminescent (panel B) and colorimetric (panel C) images. Well 1, protein standards (10 to 250 kD); wells 2 to 5, isolated nuclei; well 6, cells; well 7, cell lysates; wells 8 to 9, isolated nuclei. The wells loaded with nuclei samples correspond to independent isolations.

**Fig S10.**
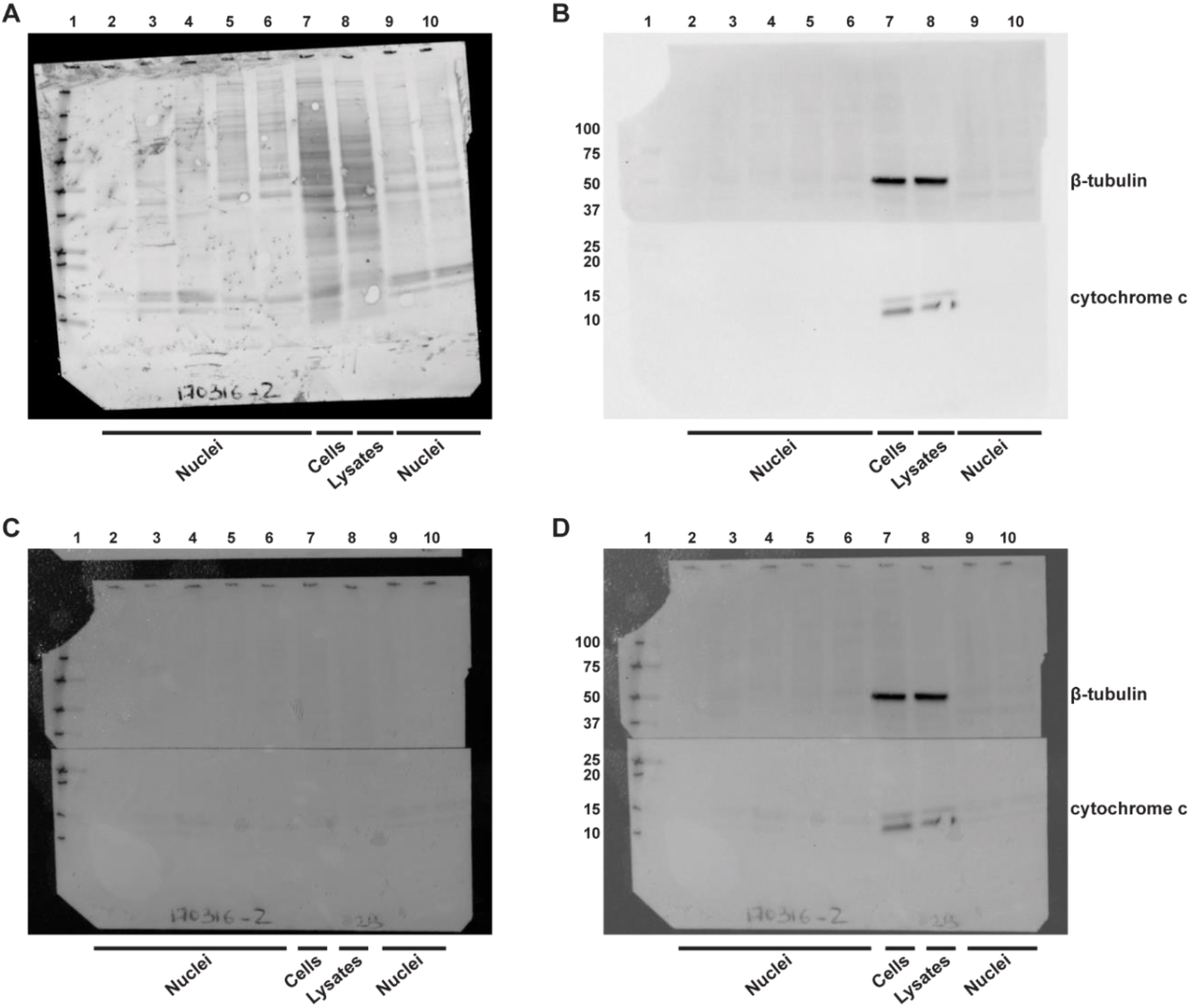
Immunoblot analysis of isolated nuclei, cells and whole cell lysates for the cytoplasmic protein β-tubulin and the mitochondrial shuttling protein cytochrome c (cyt c) (related to Fig. 1B). (A) Amido black staining of the membrane for detection of total proteins. (B) The membrane was cut in two pieces; the upper part was incubated with the antibody specific to β-tubulin and the lower part for cytochrome c. The numbers on the left side of the panel correspond to the molecular weight (kD) of the protein standards. (C) Colorimetric image of the membrane. (D) Overlay of the chemiluminescent (panel B) and colorimetric (panel C) images. Well 1, protein standards (10 to 250 kD); wells 2 to 5, isolated nuclei; well 6, cells; well 7, cell lysates; wells 8 to 9, isolated nuclei. The wells loaded with nuclei samples correspond to independent isolations.

## Legends for Tables S1 to S10

**Table S1.** Ratio of mass isotopologues of each metabolite in isolated nuclei incubated with [U-^13^C]substrates. The different mass isotopologues per metabolite (column A) are indicated by “Mn” where n corresponds to the number of ^13^C-carbons (see also Table S6).

**Table S2.** Ratio of mass isotopologues of each metabolite in whole cell lysates incubated with [U-^13^C]substrates. The different mass isotopologues per metabolite (column A) are indicated by “Mn” where n corresponds to the number of ^13^C-carbons (see also Table S6).

**Table S3.** Overview of the putative interactors associated with the bait enzymes of TCA cycle. The table is related to Fig. 3A. Columns A to C correspond to different identifiers for a biotinylated protein; NCBI Entrez Gene (A), UniProt ID (B) and Protein Name as per UniProt (C). Column D specifies the comparison between a cell line expressing a bait enzyme and the parental *HeLa* cells. The cells expressing the bait enzyme are represented with the NCBI Entrez Gene symbol of the corresponding bait enzyme followed by an *in-house* ID which is given in parenthesis (e.g. cells expressing the PDHB fused with the biotin ligase and a FLAG tag are given as PDHB(C027)). Note that, N-IDH2(N544) refers to the case of IDH2 where the biotin ligase and the FLAG tag were fused at the N-terminus of the enzyme. Columns E to I report a number of summary statistics (limma analysis) for the biotinylated proteins (columns A to C) and the indicated comparisons (column D). Column E refers to the average (across all biological replicates) log2-fold change of the abundance of the biotinylated proteins in the cells expressing a bait enzyme relative to the parental *HeLa* cells. Column F corresponds to the average log2 abundance level of each biotinylated protein across all samples. Column G is the moderated *t*-statistic. Columns H and I show the associated *p* value and the false discovery rate (fdr) adjustment for *p* values using the Benjamini and Hochberg’s method, respectively. Column J reports the bait enzyme(s) with which a biotinylated protein is significantly associated. Column K refers to Fig. 3A and specifies the cluster a biotinylated protein belongs to. Columns L and M report the number of MS runs and biological replicates, respectively, each biotinylated protein was detected. Column N shows the number of quantified unique peptide matches per biotinylated protein.

**Table S4.** Gene ontology enrichment analysis for the clusters of correlating putative interactors across all bait enzymes of TCA cycle. The table is related to Fig. 3A. Each cluster (column A) was analyzed with g:Profiler for gene ontology enrichment for the terms depicted in column C; Biological Process (GO:BP), Cellular Component (GO:CC) and Molecular Function (GO:MF). The columns are a direct output from g:Profiler.

**Table S5.** Subcellular localization of the top putative interactors of each bait enzyme of TCA cycle. The table is related to Fig. 3B. Column A shows the bait enzyme defined by the NCBI Entrez Gene symbol. Columns B to D show different identifiers for the biotinylated proteins that were significantly enriched for each bait enzyme; NCBI Entrez Gene (B), UniProt ID (C) and Protein Name as per UniProt (D). Columns E to G were retrieved from UniProt database; Column E reports the UniProt’s manual assignment of subcellular localization for the reviewed human genes corresponding to the respective biotinylated proteins. Columns H to I report the subcellular localization for each biotinylated protein as retrieved from Gene Ontology. Columns J to V refer to localization evidence from Human Protein Atlas. Column W provides the merged localization data from all previous databases. Column X reports the final localization category assigned to each biotinylated protein based on the information provided in Column W. “Nucleus” and “Mitochondria” refer to proteins only detected in the nucleus and mitochondria, respectively. “Nucleus shared” refers to proteins that are detected in the nucleus and any other compartment. This category includes also the proteins that are detected in the nucleus and mitochondria. “Mitochondria shared” refers to proteins that are detected in the mitochondria and any other compartment excluding the nucleus. “Other” includes the proteins that are detected in any compartment other than the nucleus or the mitochondria. This column was utilized for the generation of Fig. 3B

**Table S6.** Ion fragments for the quantification of the metabolites with GC-MS. Column B refers to the metabolite following the chemical derivatization with methoxyamine hydrochloride and *N*-methyl-*N*-(trimethylsilyl)trifluoroacetamide.

**Table S7.** Destination vectors and entry clones for the generation of the engineered bait enzymes fused with the biotin ligase and a FLAG tag.

**Table S8**. Primers for the generation of the engineered constructs and oligos.

**Table S9.** Sequencing primers for the engineered constructs of the bait enzymes fused with the biotin ligase and a FLAG tag.

**Table S10.** Gene ontology enrichment analysis of the putative interactors of IDH2 carrying the biotin ligase at the N-terminus. The columns are a direct output from g:Profiler which was used for the gene ontology enrichment for the terms noted in column C; “Biological Process” (GO: BP), “Molecular Function” (GO: MF) and “Cellular Component” (GO: CC).

